# A kinetic basis for curvature sensing by septins

**DOI:** 10.1101/2022.05.16.492121

**Authors:** Wenzheng Shi, Kevin S. Cannon, Brandy N. Curtis, Christopher Edelmaier, Amy S. Gladfelter, Ehssan Nazockdast

## Abstract

The ability of cells to sense and communicate their shape is central to many of their functions. Much is known about how cells generate complex shapes, yet how they sense and respond to geometric cues remains poorly understood. Septins are GTP-binding proteins that localize to sites of micron-scale membrane curvature. Assembly of septins is a multi-step and multi-scale process but it is unknown how these discrete steps lead to curvature sensing. Here we experimentally examine the time-dependent binding of septins at different curvatures and septin bulk concentrations. These experiments unexpectedly indicated that the curvature preference of septins is not absolute but rather is sensitive to the combinations of membrane curvatures present in a reaction, suggesting there is competition between different curvatures for septin binding. To understand the basis of this result, we developed a kinetic model that connects septins’ self-assembly and curvature sensing properties. Our experimental and modeling results are consistent with curvature-sensitive assembly being driven by cooperative associations of septin oligomers in solution with the bound septins. When combined, the work indicates septin curvature sensing is kinetically determined, sensitive to bulk concentration, and the available membrane curvatures. While much geometry-sensitive assembly in biology is thought to be guided by intrinsic material properties of molecules, this is an important example of how kinetics can drive mesoscale curvature-sensitive assembly of polymers.

**Significance Statement:** Cells use their membrane curvature to coordinate the activation and spatiotemporal compartmentalization of molecules during key cellular processes. Recent works have identified different proteins that can sense or induce membrane curvature from nano- to micron-scale. Septins are nanoscopic cytoskeletal proteins that preferentially bind to membranes with a narrow range of micron-scale curvatures. Yet the sensing mechanism remains ambiguous. Using a combination of microscopy and kinetic modeling, we show that, unlike most proteins that sense curvature in a single protein scale, curvature sensing in septins is determined kinetically through their multi-step hierarchical assembly on the membrane. This introduces a novel kinetic basis of fidelity, where the same protein can be deployed for differential binding sensitivities in different cellular contexts.

---

Cells and their internal compartments take on a spectacular array of shapes. How cells generate and utilize their shape to best suit their functions has been a central question in biology since the advent of the light microscope. Since then, much has been learned about how cells generate complex shapes, though little is known about how they sense their shape. Most of the work done on cell shape sensing has been done in the context of endocytosis and vesicle trafficking, where nanometer-scaled proteins assemble onto nanometer-scaled membrane curvatures. For example, proteins containing BAR domains and/or amphipathic helices are well known for their roles in both sensing extreme curvature and deforming the membrane to aid in vesicle formation (1–4). A combination of electrostatic and hydrophobic interactions allow for high-affinity binding of these proteins to curved membranes (1, 4). Convexly (or positively) curved membranes are characterized by an increase in the size and number of lipid packing defects, or exposed hydrophobic regions between lipid species, which act as binding sites for amphipathic helices (5, 6). Cells can use the complex physicochemical landscape of membrane curvature, which includes the shape and size of lipid head groups, the length of fatty acyl chains, and the electrostatic content as important features to drive preferential assembly of proteins onto curved surfaces (7–9).

Changes to cell shape, like those that occur during cytokinesis or polarized growth, require changes on a micrometer scale. This raises a fundamental question: How can cells use nanometer-scaled assemblies to detect micrometer-scale changes in membrane curvature? This disconnect in scale between molecules and geometry is striking, as a micrometer-scale curvature is essentially a flat surface for the typical peripheral membrane protein. However, these shallow curvatures are found throughout the cell at both organelles and the plasma membrane, suggesting cells must have the means to detect structures of this magnitude. SpoVM, a bacterial protein involved in sporulation, is one of the few documented micron-scale curvature sensors and uses a unique amphipathic helix to embed into the membrane similarly to nanometer-scale curvature sensors (10). Another documented class of curvature sensors in eukaryotes are septins.

Septins are GTP-binding proteins that localize to sites of micrometer-scale membrane curvature both in yeast through humans (9, 11, 12). At these sites, septins coordinate cell cycle progression, cytokinesis, and aid in the defense against invading microorganisms in host cells (13–15). Septin malfunction is implicated in various cancers and neurodegenerative diseases (16). The four essential mitotic septin polypeptides from *Saccharomyces cerevisiae*, Cdc3, Cdc10, Cdc11, and Cdc12, are assembled into palindromic, rod-shaped hetero-oligomeric complexes that constitute the minimal septin unit. Septin assembly at sites of membrane curvature begins with the binding of the septin oligomer complexes to the membrane, whose curvature sensitivity is also mediated by an amphipathic helix (9, 17). Membrane-bound oligomers will then diffuse and interact with one another to form filaments in a process termed diffusion-driven annealing (18). Septin filaments can either fragment or connect with nearby filaments to form higher-order structures including pairs, rings, and gauzes (19–21). The mechanisms by which these hierarchical steps of assembly contribute to septin’s curvature sensing function remain largely unexplored.

In earlier studies we used mixtures of beads of multiple diameters (poly-dispersed) to study the curvature-sensitive binding of septins and found that septin adsorption is maximized on 1 *μ*m beads, consistent with the curvature where they are found to localize in cells (9, 12). As we will show here, in experiments where there is a single curvature, i.e. bead size, present in the assay (mono-dispersed case) or two curvatures in the assay (bi-dispersed case) lead to different adsorption patterns and apparent curvature sensitivities. These observations suggest that beads of different curvatures can interact/compete non-locally and that the “optimal” curvature sensed by septins is not fixed - it is instead a function of the assembly process and available membrane platforms.

Here we use a combination of single molecule imaging, scanning electron microscopy, time-lapse confocal microscopy, biophysical modeling, and computer simulations to propose a unified kinetic model that connects several different time-dependent processes to the observed curvature sensing of septin assemblies. Our model correctly predicts the observed time-dependent binding of septins at different bulk concentrations to beads of different radii. It also predicts the different behaviors observed in mono- and bi-dispersed assays. From our model, the difference between the assays arises due to the competition between beads of different radii as a shared, finite pool of septins is depleted by curvature-dependent kinetic processes. We provide experimental support of this prediction by varying the total membrane surface area to alter the degree of depletion. We find that processes of diffusion and end-on annealing of bound filaments are not sufficient for curvature sensing; instead we hypothesize that they control the rate of formation of a foundation layer of septins. The model predicts that septin binding is dominated by cooperative interactions between the bound septin filaments in that foundation layer and free septin oligomers in the bulk to create curvature-dependent assemblies.

## Results

### Time-dependent adsorption of septins

To study the assembly process of membrane-bound septins, we used confocal microscopy and measured time-dependent septin adsorption onto membrane-coated beads with assays that were developed and used in our earlier studies (9, 22). One main difference between our current and past studies is that here the adsorption is measured over time, whereas in the previous studies we only reported the adsorption at steady-state. Secondly, here we directly compare assays where all spherical supported lipid bilayers are the same radius (mono-dispersed) with assays of two different radii (bi-dispersed), also referred to as the *competition case*. As in previous studies, total membrane surface area is kept constant between all conditions unless specifically stated, and the surface area of 1 *μ*m to 5 *μ*m beads in bi-dispersed cases is always 1:1.

The fluorescence intensity of septins was measured through time and as a function of bead size with diameter 1 *μ*m or 5 *μ*m, each at bulk concentrations 6.25, 12.5, 25 and 50 nM septins in mono-dispersed assays (Fig. 1A-B). The time-dependent adsorption onto 1 *μ*m and 5 *μ*m beads in competition (bi-dispersed assays) at bulk concentration 25 nM is displayed in Fig. 1C. The main observations can be summarized as follows:

1. The time-dependent adsorption resembles a sigmoidal curve and can be divided into three sequential periods, regardless of the bead radius and bulk concentration. (i) Initiation period, where the adsorption remains very limited and increases slowly, (ii) growth period, where both the adsorption and adsorption rates increase superlinearly with time, and (iii) saturation period, where the adsorption rate begins to slow down and adsorption approaches its steady-state value (Fig. 1D); also, see section F in the supplementary materials.
2. In all mono-dispersed assays, the initiation and growth timescales are reduced (i.e. adsorption rates are increased) by increasing the bulk concentration of septins as well as membrane curvature, which is inversely proportional to the beads’ radii.
3. Most interestingly, we observe a larger steady-state adsorption on 1 *μ*m compared to 5 *μ*m beads only when the two bead sizes are co-incubated. In contrast, when only a single bead size is present in the reaction, we find that the steady-state adsorption levels on 5 *μ*m beads are larger than those on 1 *μ*m beads while controlling for surface area and concentration of septins in the bulk (Fig. 1C and 1F). Similar steady state differences were seen when comparing 1 *μ*m and 3 *μ*m beads in previous studies (9). Notably higher adsorption on 1 *μ*m beads in the competition setting is associated with its shorter lag times and faster growth rates, compared to 5 *μ*m beads, indicating kinetic differences in the assembly process on the two curvatures when they coexist.

**Fig. 1.**
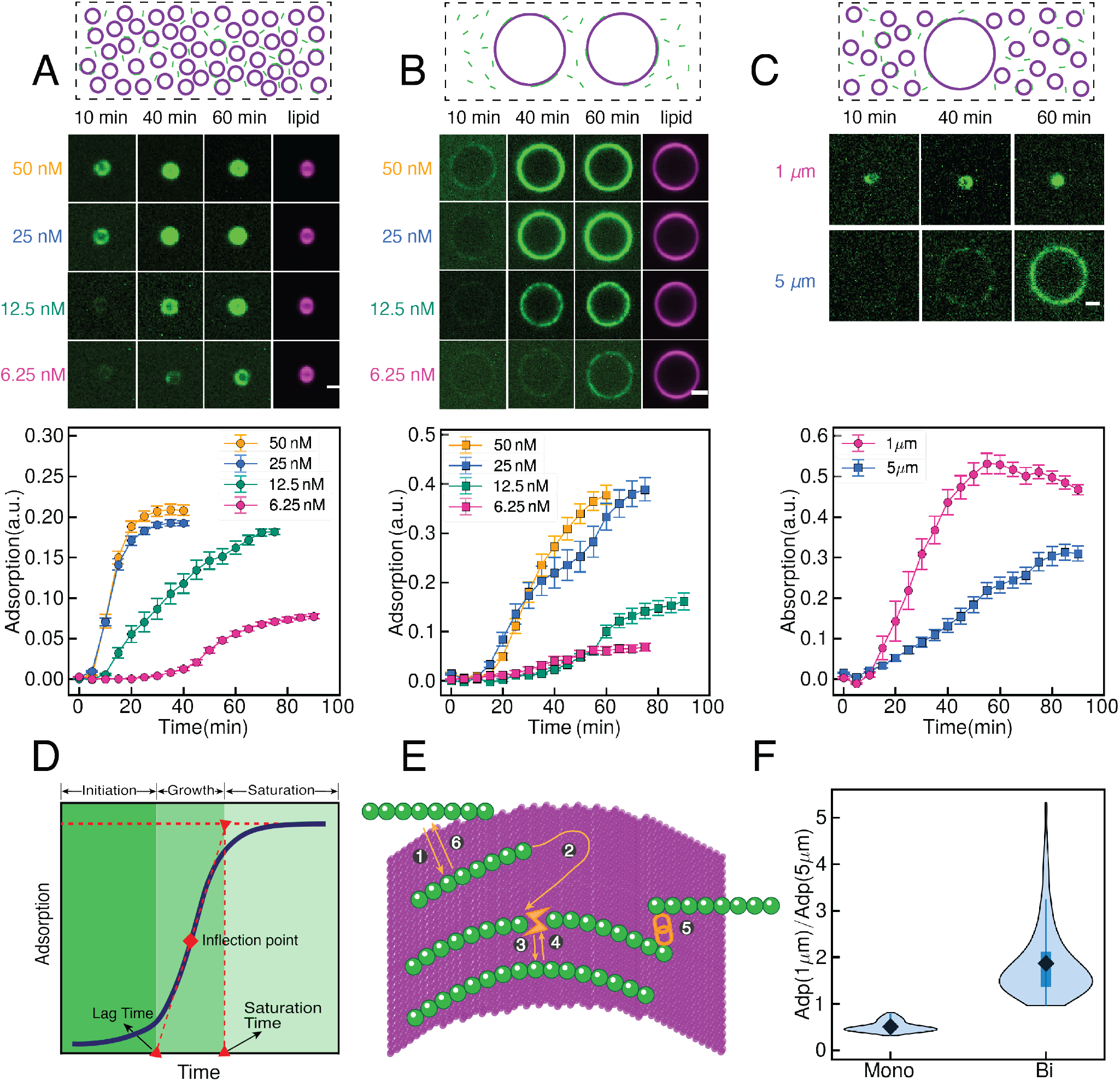
Time-lapse microscopy on supported lipid bilayers shows context-dependent septin assembly kinetics. (A-C) First row: schematic representations of the view of mono-dispersed (A and B) and bi-dispersed assays (C); the total membrane surface area is kept fixed in all experiments. Second row: The time variations of the representative focal slices of septin adsorption (green) at various concentrations onto (A) mono-dispersed 1 *μ*m, and (B) 5 *μ*m beads; and (C) bi-dispersed mixture of beads at bulk concentration *n_b_* = 12.5 nM. The lipid channel is in magenta, and the scale bars are 1 *μ*m in (A), 2 *μ*m in (B) and 1 *μ*m in (C). Third row: quantification of septin adsorption. Error-bars correspond to the standard error. (D) Diagram of the adsorption process, separated into three regimes: initiation, growth and saturation. (E) Sub-processes involved in the septin adsorption process: ① binding of a single septin oligomer in the bulk; ② the diffusion of bound septins on the membrane; ③ their polymerization through end-on annealing; ④ fragmentation of bound septin filaments into shorter ones; ⑤ cooperative binding of bulk septins, and ⑥ unbinding. (F) The ratio of septin adsorption at steady state on 1 *μ*m beads to septin adsorption on 5 *μ*m beads for mono- and bi-dispersed assays.

These results indicate that maximum septin adsorption to different curvatures is not absolute, but can vary depending on the presentation of single or multiple curvatures in the same reaction. The time-dependent adsorption of septins is determined by the summation of different processes involved in septin assembly including membrane binding/unbinding, diffusion, end-on annealing into filaments, fragmentation of filaments, and potentially binding/unbinding of septin-septin interactions that may form layers or lateral assemblies (Fig. 1E). Which of these many steps, if not all, drives the curvature-dependent assembly of septins in these different experimental regimes? We combine these experiments with modeling to examine the relationship between membrane curvature and bulk concentration with each of these discrete steps and their potential impact on determining initiation, growth and saturation periods. Note that the adsorption process typically takes tens of minutes, which is thousands of times longer than the time-scale associated with a single oligomer’s dwell time (9). This long time interval makes using time-costly atomistic dynamic simulations and its more coarse-grained variations incompatible with the questions we seek to answer; thus, we use a Smoluchowski reaction kinetic framework (23, 24), informed by the findings of this work and previous ones, to simulate the time-dependent assembly of membrane-bound septins.

### Binding/unbinding kinetics of single septin complexes

Septin assembly begins with single septin oligomers binding the membrane from the bulk. Our earlier studies show that the binding rate of a single septin oligomer monotonically increases with curvature (9). Curvature-dependent binding rates have also been reported for the only other characterized micrometer-scale curvature sensor, SpoVM (10). Based on these findings, we hypothesized that curvature-dependent binding of septins might be the main driver for the assembly and curvature sensing of septin assemblies. To test this hypothesis and gain measured parameters for the kinetic model, we used a previously developed assay (9), where solutions of recombinantly purified, non-polymerizable septin oligomers are mixed with membrane-coated beads of different curvatures (radii). Here we expand the range of bulk concentrations and bead radii used in (9). The number and rate of binding events as well as the dwell time for single oligomer were measured using near-TIRF microscopy on beads of different sizes (Fig. 2A). As previously shown for different bead sizes, binding rates change as a function of curvature, however these lead to a monotonic increase in binding rate rather than actual curvature sensitivity, suggesting the binding rate on its own is insufficient to generate a curvature optimum (Fig. 2B).

**Fig. 2.**
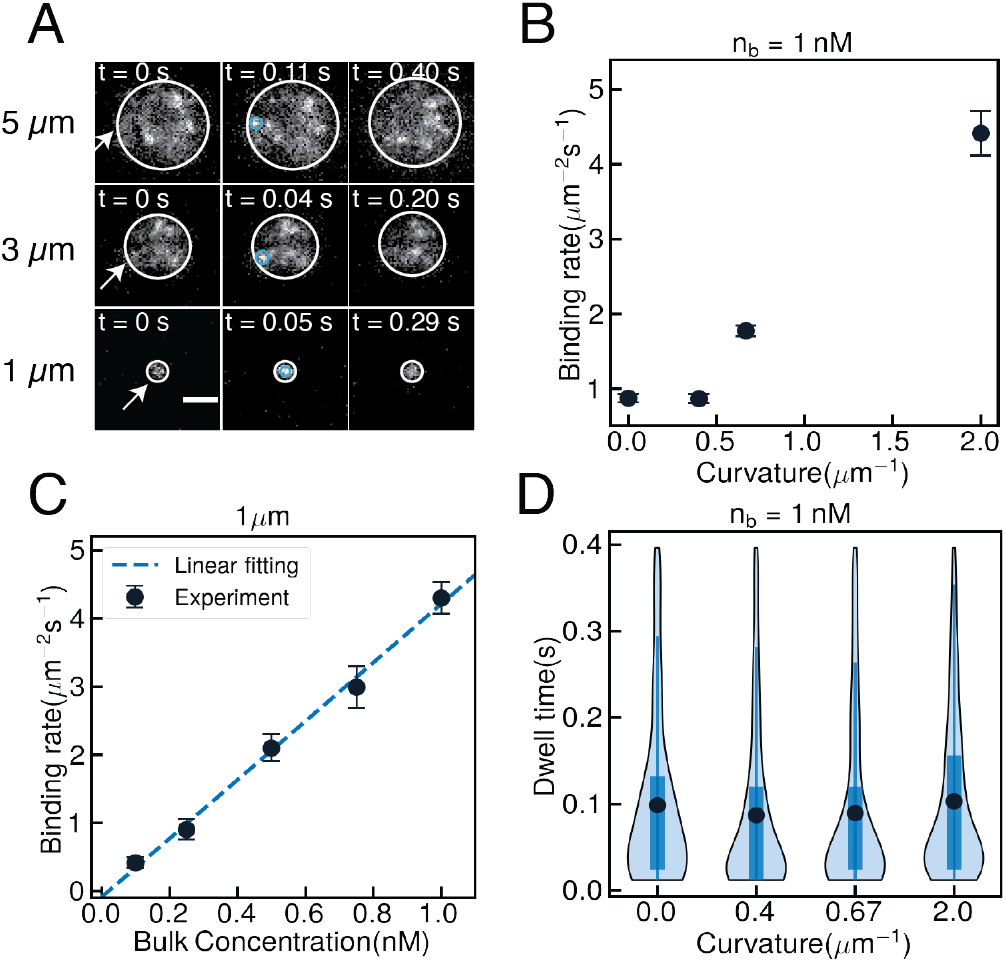
Single septin complexes can only detect changes in membrane curvature through their binding rate. (A) Representative near-TIRF images showing single oligomers binding and unbinding events on various membrane curvatures. Scale bar is 2 *μ*m. (B) Measured binding rates (*J*_on_) for a single oligomer binding onto various membrane curvatures, *κ* = 1/*r*, at 1 nM bulk concentration. *N* > 200 events for each curvature. Error bars represent the standard error. (C) Measured binding rates of single septin oligomer onto membrane curvature *κ* = 2 *μ*m^−1^ at various septin concentrations. N > 200 events for each concentration. Error bars represent the standard error. (D) Violin plots highlighting the distribution of measured dwell times for a single septin oligomer on different membrane curvatures at 1 nM bulk concentration. N > 200 events per curvature. Filled circles show the experimentally measured mean values.

We incorporated the binding rates into the kinetic model with considerations for what is known about the molecular requirements of curvature sensitivity of septins via conserved amphipathic helices (AH). Other studies of curvature-sensing proteins suggest that membrane binding occurs in part through AH insertion into lipid packing defects, which are induced through curvature and thermal fluctuations within the membrane (6, 10, 25). Intercalation of AH domains into the defects is driven by attractive electrostatic interactions between the AH domains and the membrane (5, 26).

Following these findings, we model the adsorption/desorption process by dividing it into three sub-processes: (i) the formation and healing of lipid packing defects; (ii) the binding of the single septin to a given defect; and (iii) the unbinding of the membrane-bound oligomer to the bulk. Following this, the kinetic equations describing septin oligomer’s adsorption with time, *n^s^*, are expressed as

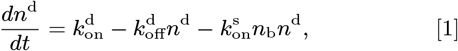

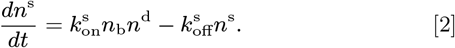

where superscript d, s, respectively, refer to defects, membranebound septins, superscript b refers to septins in the bulk, and letter *n* denotes surface density for *n*^s^, and *n*^d^ and bulk number density for *n*^b^. The likelihood of having larger and longer-lived defects with higher membrane curvature is captured by an increase in the rate of defect formation, 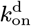, leading to monotonically increasing binding rates with curvature. This is in line with the experimental observations shown in Fig. 2B.

Within our model, the rate of septin binding per unit area (*J*_on_) is the first term on the right-hand-side of Eq. 2: 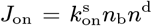. At steady-state (*dn*^d^/*dt* = 0, *dn*^s^/*dt* = 0), *J*_on_ is given by

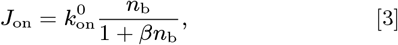

where 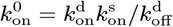 and 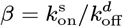. When the binding rate of oligomers to defects is significantly smaller than the healing rate of defects, 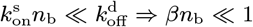, the model predicts a linear increase of septin binding rate with bulk concentration, which is what we observe experimentally (Fig. 2C) at small bulk concentration *n*_b_ ≤ 1 nM. At sufficiently large bulk concentrations 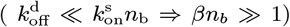, the net binding rate becomes independent of bulk concentration 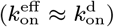. This is consistent with the observations in Fig. 1A,B that show time-dependent adsorption curves at 25 nM and 50 nM are nearly identical for any given bead size.

Next, we expanded on previous work to experimentally determine the dwell times of bound septins on a range of membrane curvatures at low septin concentration (Fig. 2D). The unbinding rate of a single septin oligomer, 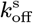, is defined as the inverse of their average dwell time. As shown, the measured dwell times, and thus 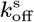, remain mostly unchanged with variations of curvature. This observation is consistent with the idea that, upon septin insertion, the septin and membrane molecules locally reconfigure to an energy state that is independent of the membrane’s macroscopic energy/curvature; this, then, leads to nearly curvature-independent unbinding rates.

Setting *dn*^s,d^/*dt* = 0, the steady-state adsorption of single oligomers is given by the ratio 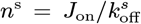. Given that our measurements show a monotonic increase of binding rate with curvature (*J*_on_ ↑ as *κ* ↑) while 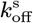 remains roughly constant (*k*_off_ = constant as *κ* ↑), the absorption should increase monotonically with curvature (*κ* ↑ leading to *n*^s^ ↑). Thus, single septin binding/unbinding kinetics cannot explain the observed curvature preference of septins. On the other hand, these observations together with the results of time-dependent adsorption on mono- and bi-dispersed assays support the idea that the preferred curvature for septin binding to membranes is determined kinetically, raising the question: what additional steps in assembly may be controlling the assembly rates? We proceed now with the development of the kinetic model to include multiple steps of assembly along with the experimental measurement of parameters for each step.

### Diffusive motion of septins on the membrane

Once a septin oligomer is bound to the membrane, it undergoes diffusive motion and interacts with nearby oligomers/filaments to form longer filaments in a process referred to as *diffusion-driven annealing* (Fig. 1E, 2-4). This is a key step in determining the organization of septin filaments on membranes. We hypothesized that differences in diffusion on different curvatures could impact elongation/filament lengths. Using our single molecule TIRF assay and particle tracking software (27) we tracked the position of bound single non-polymerizable septin oligomers on curved and flat membranes (Fig. 3A). We were only able to obtain the septins’ trajectories on beads of diameter 3 *μ*m, 5 *μ*m and not for beads of diameter 1 *μ*m, because these beads were too small for sufficient track lengths in the field of view which was limited by the height of the evanescent wave of the TIRF imaging. Fig. 3B-D show the mean squared displacements (MSD) computed from particles’ trajectories in log-log scale, and compare them against the MSD for single septins on planar membranes. We find that the single septins undergo nearly diffusive motion on a planar membrane (MSD ~ *t^α^*, where *α* ≈ 1 for purely diffusive motions), consistent with membrane responding as a Newtonian fluid in tangential directions. Interestingly, we also find that single septins exhibit sub-diffusive motion (*α* < 1) on curved membranes, ultimately reaching to a nearly constant MSD at long times, which is indicative of an elastic response at long time, as shown in Fig. 3C-D. Thus, diffusion of single septin complexes qualitatively changes on flat versus curved membranes, raising the possibility of this being a source of kinetic difference relevant for curvature sensitivity.

**Fig. 3.**
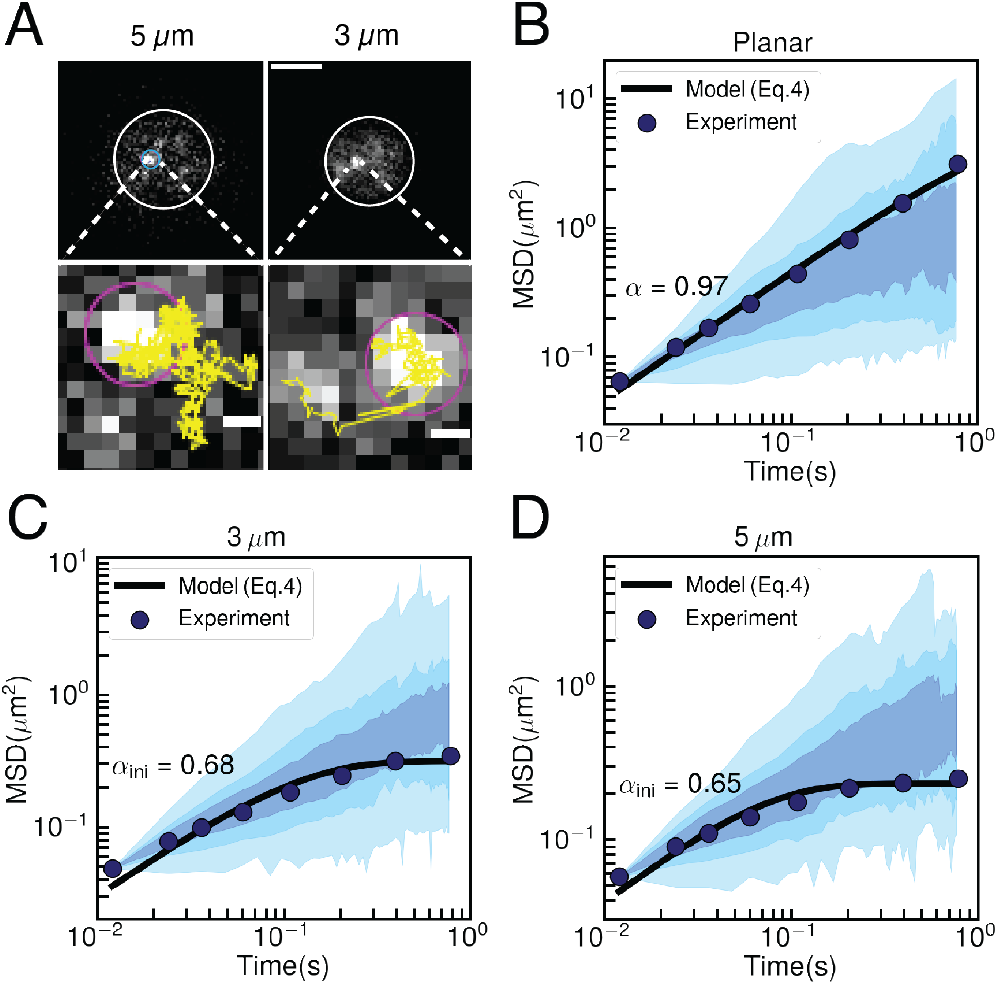
Single septin complexes display sub-diffusive behavior on curved membranes. (A) Top panel: Representative near-TIRF images of single septin complexes on membrane-coated beads of radii 5 *μ*m and 3 *μ*m, respectively. The Scale bar is 0.25 *μ*m. Bottom panel: Individual particle tracks (yellow lines) for beads of radii 5 *μ*m and 3 *μ*m, respectively. The scale bar corresponds to 0.25 *μ*m. (B-D) Mean squared displacements (MSDs) of single septin oligomers vs time on a flat membrane (B), and membrane-coated beads of radii 3 *μ*m (C), and 5 *μ*m (D). The filled circular symbols denote the mean values computed from analysing the trajectory of more than 500 particles, and the solid lines represent the fit from Eq. 4 with *N* = 1. Shaded areas represent spread in experimental data in the interquartile range (darkest, 25% – 75%), interdecile range (next darkest, 10% – 90%), and full range (lightest, 0% – 100%).

**Fig. 4.**
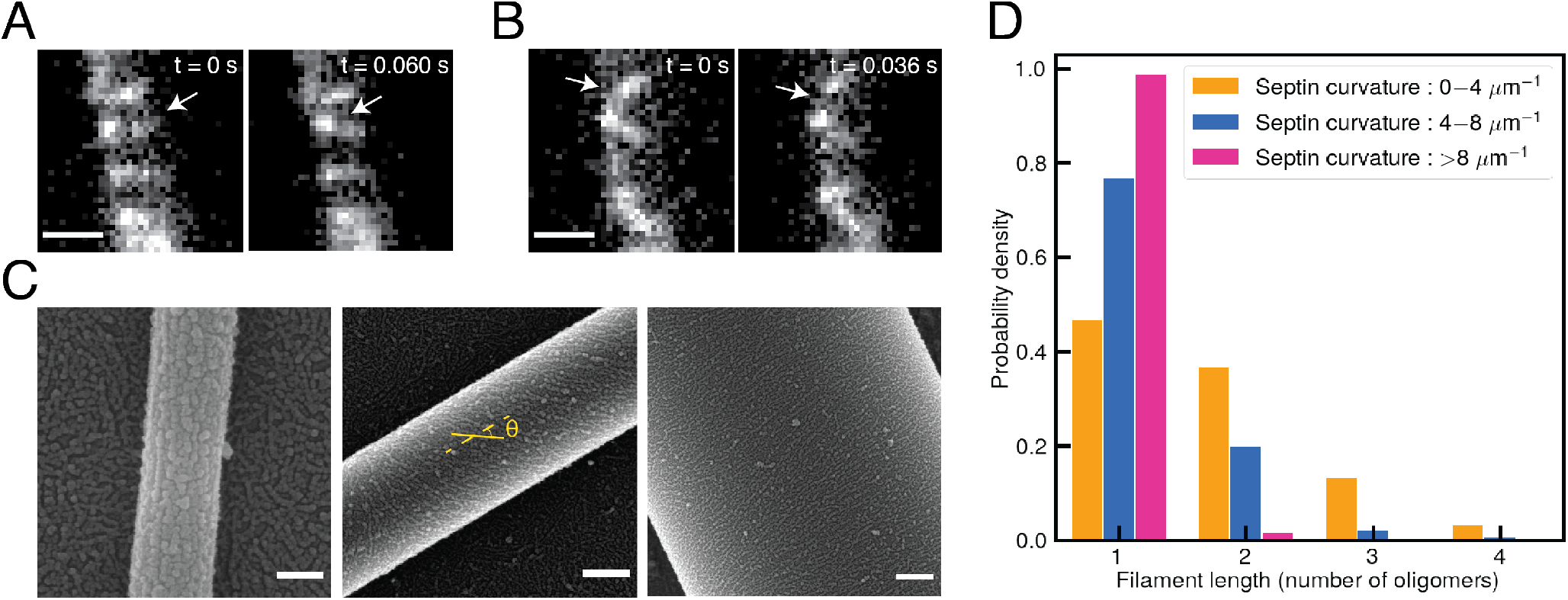
The length distribution of septin filaments is curvature-dependent. (A-B) Near-TIRF images of septin filament end-on annealing (A) and fragmentation (B) on membrane-coated rods of diameters 46 – 1508 nm. Scale bar is 1 *μ*m. (C) SEM images of septin filaments on rods of different diameters. One of the septins is label as a yellow solid line and the yellow dashed line is the long axis of the rod. The angle between septin axis and the rod axis is defined as *θ*. Scale bars are 0.1 *μ*m. (D) Probability density of septins’ lenght (number of oligomers per septin) at different septin curvature intervals. The curvature is defined as *κ*^s^ = (*R* sin *θ*)^−1^ where *R* is the radius of the membrane-coated rod and *θ* is the angle of the septin filament with respect to the long axis of the rod (zero when septin filament lies along the long axis and *π*/2 when septin filament lies orthogonal to the long axis).

We use this data to incorporate membrane diffusion into the kinetic model. As discussed previously, AH binding to the membrane is driven by electrostatic interactions between the polar residues of the AH and lipid head groups followed by incorporation of nonpolar AH residues into the hydrophobic core of the membrane. These attractive interactions can be modeled, in their simplest form, as spring-like connections between the membrane and a bound septin. These connections lead to an elastic response. Here, we use a phenomenological model to describe the MSD of bound septins. Specifically, we use a Kelvin-Voigt (a parallel spring and dashpot) model to describe the displacements of septins in response to thermal forces; see Fig.S1. The dashpot element models the viscous resistance to a septin’s motion on the fluid membrane, and the spring element models the attractive interactions between septin’s AH domain and the membrane. The resulting overdamped Langevin equation describing the septin dynamics is *γ*d**x**/d*t* = **F**^Br^ – *K*_sp_**x**, where **F**^Br^ is the fluctuating thermal force, *γ* is the drag coefficient of the septin moving through the membrane, and *K*_sp_ is the spring coefficient modeling the attractive interactions between AH domain and the membrane.

Given that in our micron-bead assays and in living cells, the average length of septin filaments are most likely significantly longer than a single oligomer, we need to know how septin’s sub-diffusive motion is changed with its length. Since we cannot experimentally control the length distribution of septin filaments throughout the adsorption process, we resort to modeling to describe the MSD of septins of different lengths. To extend the Langevin equation to septins made of *N* oligomers we must specify how septin’s drag coefficient, *γ*, and spring coefficient, *K*_sp_ change with *N* (length). The drag coefficient of a rod-like inclusion on a spherical membrane is a complex function of its length, membrane viscosity, viscosity of surrounding bulk fluid, and the sphere radius (28). Using scaling analysis, presented in section A of the supplementary materials, we show that the drag of a rod-like particle along its axis scales linearly with its length in membranes coated on spherical rigid-like beads. We assume that the effective spring coefficient scales linearly with the number of AH-membrane interactions, which also scales linearly with the septin length. Implementing all these assumptions and integrating the Langevin equation, the MSD for a septin made of *N* oligomers is

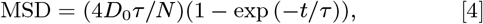

where *τ* = *γ/K*_sp_, and *D*_0_ = *k*_b_T*/γ* is the diffusion of a single oligomer without AH-membrane interactions. Note that MSD scales inversely with the length of septins (MSD ~ 1/*N*). The details of the model are provided in section A of the supplementary materials. The results of fitting this model on the experimental data for single oligomers (*N* =1) are in very good agreement and are shown as solid lines in Figs. 3B-D.

Our model can also be used to explain the observed difference of MSD on flat and curved membranes. Recall that our TIRF microscopy results show that increasing the membrane curvature increases septin’s binding rate (Fig 2B). Previous work ascribed the curvature preference of AH domains to the increasing size and density of lipid packing defects with increasing curvature (5, 29). Thus, we model defect density, or number of AH binding sites, as proportional to the spring coefficient, *K*_sp_, which increases with curvature in our model. Assuming the drag coefficient, *γ*, is not a strong function of curvature (see Fig.S1 for the analysis that supports this claim), the increase in the spring stiffness with curvature reduces *τ* = *γ/K*_sp_ and leads to a more subdiffusive motion on curved membranes in the time window of the experiments. We compute the values of *D*_0_ and *τ* at different bead curvatures from their respective single septin MSD measurements. We then use those computed values in Eq. 4 to compute the MSD of septin filaments made of *N* oligomers at different curvatures (Fig.S1).

### Unbinding rates of septin filaments

Net septin adsorption is dependent not only on binding but also unbinding rates. Earlier in the paper we used near-TIRF microscopy (Fig. 2A) to measure the unbinding rate of single septins, and found them to be independent of curvature for a single oligomer (Fig. 2D). Next, we ask how does the unbinding rate change with septin’s length? We use a simple kinetic model to answer this question. In our model, a septin composed of *N* oligomers forms *N* attachments with the membrane. This septin is released only when all *N* attachment sites are released. We use the measured single septin oligomer unbinding rate 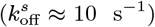 to describe the unbinding kinetics of each one of *N* attachments of a bound septin. The other key component of the model is that once an attachment is released, it can re-bind with the membrane with a rate *k*_rb_, as long as the filament remains bound through its other AH attachments. The average dwell time of a filament with *N* attachments, which is the inverse of the effective detachment rate, is computed by numerically solving a system of *N* ordinary differential equations describing the time-dependent probability of observing different attachment states. Through post-analysis of the computed detachment rate for different values of *N*, *k*_rb_ and 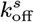 we find that the dettachment rate exponentially decreases with the number of attachment sites (length): 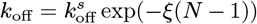, where *ξ* is an empirical function that monotonically increases with the ratio of 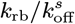. Thus, our model predicts that as the bound septins merge and form longer filaments they become exponentially more stable (longer-lived). The details of the model are provided in section B and Fig. S.2 of the supplementary materials. The value of *ξ*, and by extension *k*_rb_, is determined through the optimization process.

### End-on annealing and fragmentation processes

Filament formation is required for stable septin-membrane interactions *in vitro* and essential for septin function *in vivo* (12). As we discussed in the previous section, end-on annealing of oligomers into filaments stabilizes septins’ interaction with the membrane by dramatically extending their lifetime (reducing their unbinding rates). We define end-on annealing as the process where *two bound septins* merge and form longer filaments. We assume that end-on annealing can occur only through annealing at either end of each bound septin filament. Fig. 4A shows near-TIRF images of septin polymers prior to and after end-on annealing. Note that bound septins can also increase in length through cooperative binding of *bulk* septins to either end of the bound septin filament. Conversely, septin filaments can break into smaller filaments or individual oligomers in a process termed fragmentation, thereby reducing the dwell time on the membrane. Fig. 4B shows near-TIRF images of a septin polymer prior to and after fragmentation. For a septin filament composed of *N* single oligomers, the fragmentation can occur at (*N* – 1) points along the filaments.

Next, we examined if membrane curvature could influence the end-on annealing and fragmentation rates, and hence the length distribution of septin filaments. We turned to an assay we previously developed to examine septin filament length and curvature at nanometer resolution. This approach uses scanning electron microscopy (SEM) to image cylindrical rods of radii *R*_rod_ = 46 – 1508 nm, coated with lipids and incubated with septins (Fig. 4C). Spherical surfaces, such as micron-beads, are special in that the curvature is constant at any point and along any tangential directions. This is not true for any other surface including the cylindrical geometry of the rod, where the curvature along the filament’s axis is *κ*^s^ = 0 when the filament lies along the long axis of the rod, and changes to 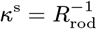, when it lies perpendicularly across the rod (along the short axis).

In our previous study we showed that the alignment angle of bound septin filaments with the main axis of the rod (shown as *θ* in Fig. 4C) is increased from nearly *θ* = 0 at *R*_rod_ =0.1 *μ*m to *θ* ≈ *π*/2 at *R*_rod_ > 0.35 *μ*m (9). Given that the curvature of a septin filament with alignment angle *θ* along its axis is *κ*^s^ = (*R*_rod_ sin *θ*)^−1^, we hypothesize that septins sense the membrane curvature in the direction of their main axis and change their alignment angle to achieve optimal curvature along their axis. This hypothesis predicts that the ensemble average curvature of septins on rods is independent of the rod’s radius. We test this prediction by post-analyzing the SEM data, which is outlined in Fig. S.5 of the supplementary materials. We find that the average curvature is indeed independent of the rod’s radius, and, thus, independent of any measure of curvature that is solely determined by rod’s geometry, such as Gaussian and mean curvatures of the membrane.

We also used the SEM data to determine the probability density distribution of the septins’ length, normalized by the length of a septin oligomer, as a function of the filament’s curvature, *κ*^s^. Fig. 4D shows the exponentially decaying length distribution of septins at different curvature intervals. At the lowest curvature interval (*κ*^s^ < 4 *μ*m^−1^), i.e. more flat, septins can form filaments that are in average made of up to four oligomers. The average length decreases as the curvature increases, and at the highest interval (*κ* > 8 *μ*m^−1^), only oligomers are present. We hypothesize that this could be due to an increased rate of fragmentation or decreased rate of end-on annealing at higher curvatures. Consistent with these observations, we constrain the ratio of fragmentation to end-on annealing rate (*k*_frag_/*k*_anneal_) to be an increasing function of curvature in our modeling of spherical membrane-coated bead assays.

### Cooperative binding

A major feature of the time-dependent assembly of septin filaments is the superlinear increase of binding with time during the growth period. In writing Eq. 2, we have assumed that the rate of defect formation and septin binding are independent of the density of bound septins. In this case, given that the unbinding rate of septins is a positive constant number, we expect the rate of adsorption (*dn*^s^/*dt*) to decrease as the adsorption (*n*^s^) is increased over time, which goes against the experimental results (Fig. 1) during the growth period. This prompted us to ask what processes can give rise to the increased slope of adsorption with time? One possible answer is that end-on annealing of bound septins produce longer filaments, thereby decreasing their unbinding rate, *k*_off_(*t*), resulting in an increase in *dn/dt* over time. This mechanism is already partially described by the combination of the length-dependent off-rate and fragmentation processes.

Alternatively, this superlinear growth may be driven by *cooperative binding* of bulk septin to the membrane, which we define as any mechanism where the bound septins facilitate the binding of additional bulk septins to the membrane. One plausible physical model for cooperative binding is that septins locally deform the membrane upon binding, which has been shown in previous computational studies (30, 31). These local deformations can increase the probability of forming sufficiently large and stable defects for recruitment of septins from the bulk, leading to a cooperative binding mechanism (6). Cooperative binding may also be produced by attractive interactions between septin AH domains and the backbones of either bound or bulk septins. Regardless of the particular physical underpinning of the cooperative binding process, from a mathematical standpoint all scenarios produce a binding rate that scales positively with the density of bound septins: *dn/dt* ~ *n*_b_*n*^s^. We model the cooperative binding kinetics of defect formation and healing in the same manner as the direct binding mechanism discussed earlier. We account for the defect formation enhanced by septin binding by changing 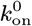 in Eq. 3 for direct binding to 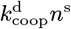 for cooperative binding.

Furthermore, Fig. 1 shows that steady-state adsorptions at the two largest bulk concentrations (*n*_b_ = 25 nM and 50 nM) remain nearly unchanged for a given bead curvature. This maximum saturation is larger in 5 *μ*m in mono-dispersed assays. This may be due to its relatively slower organization kinetics, compared to 1 *μ*m beads, allowing sufficient time for the filaments to align thereby increasing its packing efficiency. We phenomenologically model the effect of having maximum surface density by multiplying the binding rate by (1 – *n/n*^sat^), where *n*^sat^ is the maximum surface density on a particular radius. After these modifications we have:

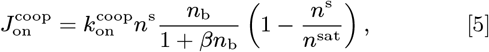

where, for simplicity, we have assumed the same value of 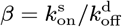.

The cooperative flux shown in Eq. 5, can either increase the length of bound septins by adding the bulk septins to either of the two free ends of bound septins (modeled as 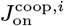 in Table 1), or it can bind single oligomers from the bulk to the membrane near pre-existing filaments (modeled as *J*^coop,1^ in Table 1). The second scenario leads to polymerization of new septins in the vicinity of pre-existing filaments.

**Table 1.**
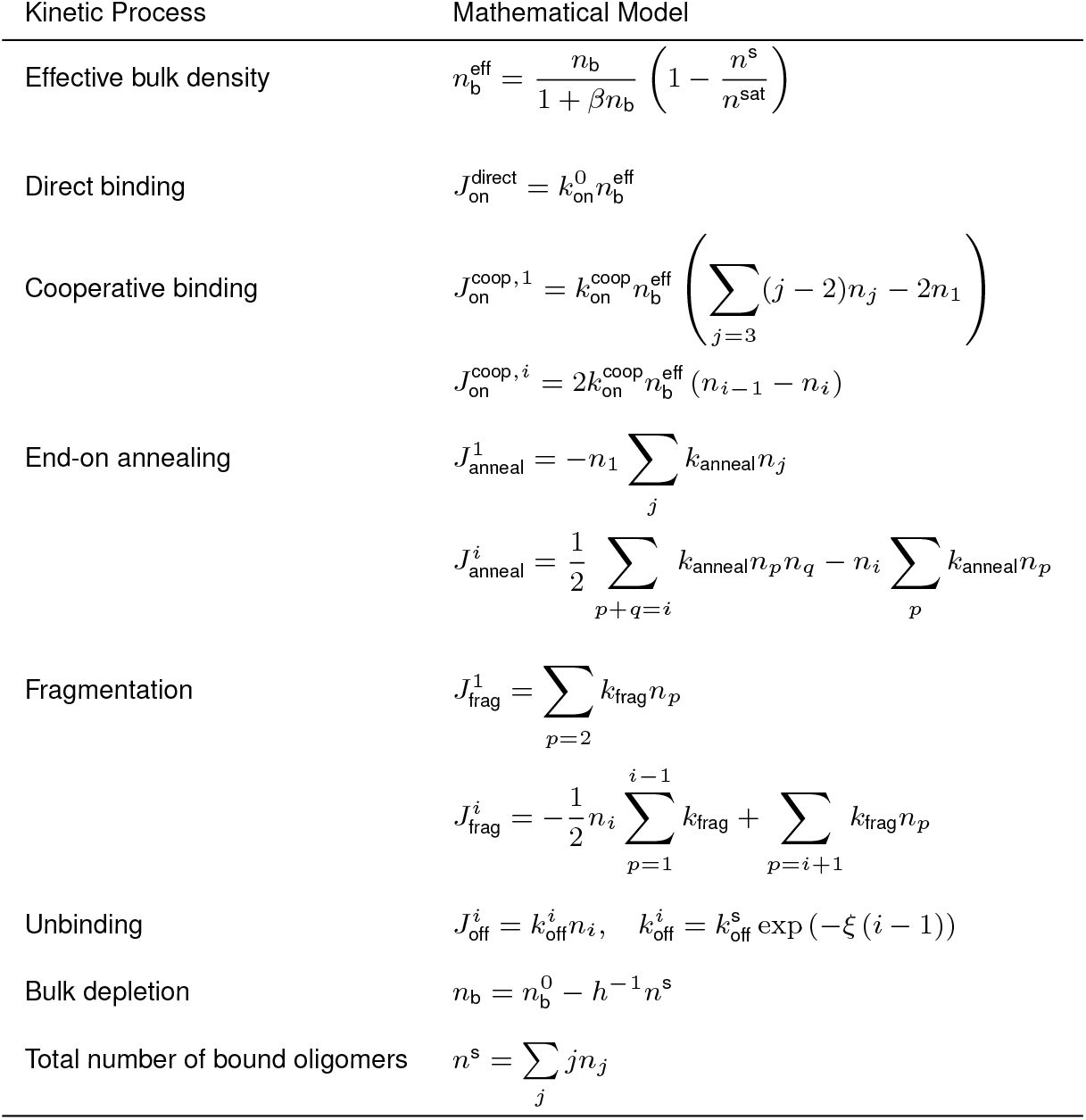
Different processes involved in septin assembly on the membrane and their associated mathematical models.

Note that all experiments are carried out in a closed system with a fixed total mass of septins, which all appear initially as oligomers in the bulk. Thus, the septins in the bulk are depleted as the adsorption is increased with time: 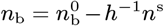. Here, 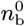 is the bulk concentration of septins at the start of the experiments, and *n*^s^ is the time-dependent surface density on each bead assay, and *h*^−1^ = *A/V* is the *experimentally known* ratio of the total membrane surface area of the beads, *A*, to the volume of the chamber containing septin oligomers in the bulk, *V*. As we shall see in the modeling section, this depletion effect must be included to predict the observed variations of adsorption with curvature on mono-dispersed vs bi-dispersed assays. Note also that in experiments involving depolymerizable single septin complexes shown in Fig. 2, the net adsorption is negligible, and depletion effects can be neglected.

Finally, we note that while the measured fluorescent intensity from confocal imaging, referred to here as adsorption, is proportional to the total mass of bound septins, it is not equal to the septin surface density: *n^s^* = Ω × Adsorption, where Ω is factor that relates adsorption with arbitrary units to surface density with the unit of number/area. In the absence of bulk depletion Ω cannot be determined independently i.e. any choice of Ω would simply re-scale the adsorption curves, and it is mathematically impossible to uniquely relate the adsorption to surface density. Including depletion effects and the observed reversal in steady-state adsorption ratio of 1 and 5 *μ*m beads in mono- and bi-dispersed assays resolves this issue and allows us to uniquely determine parameter Ω through matching the predictions against experiments and, thus, predict the total surface density of bound septins.

## Modeling and simulation methods

We have now developed the required pieces for modeling binding, unbinding, surface diffusion, end-on annealing, and fragmentation processes and their curvature dependence that ultimately determine the time-dependent adsorption and assembly of septin filaments on membranes. The equations describing these processes are summarized in Table 1. The kinetic equations describing the general assembly process are

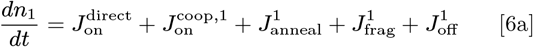

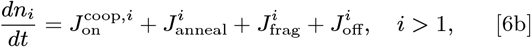

in which superscript in *J* and subscripts in *n* denote the number of oligomer units in the filament.

For filaments to polymerize, their ends must get within a critical distance of each other, which is accomplished through their sub-diffusive motion. The parameters that appear in modeling these processes are listed in Table 2. Note that the parameters labelled as “S” under source column (except for *n_i_* and *l*_ave_, which are determined by specifying the other parameters) are determined through a least-square optimization that minimizes the difference between the predicted and measured time-dependent adsorption curves at different bulk concentrations and curvatures. In order to solve this inverse problem, we first need a method to solve the direct problem: given all the parameters in Table 2, predict the time-dependent adsorption.

**Table 2.**
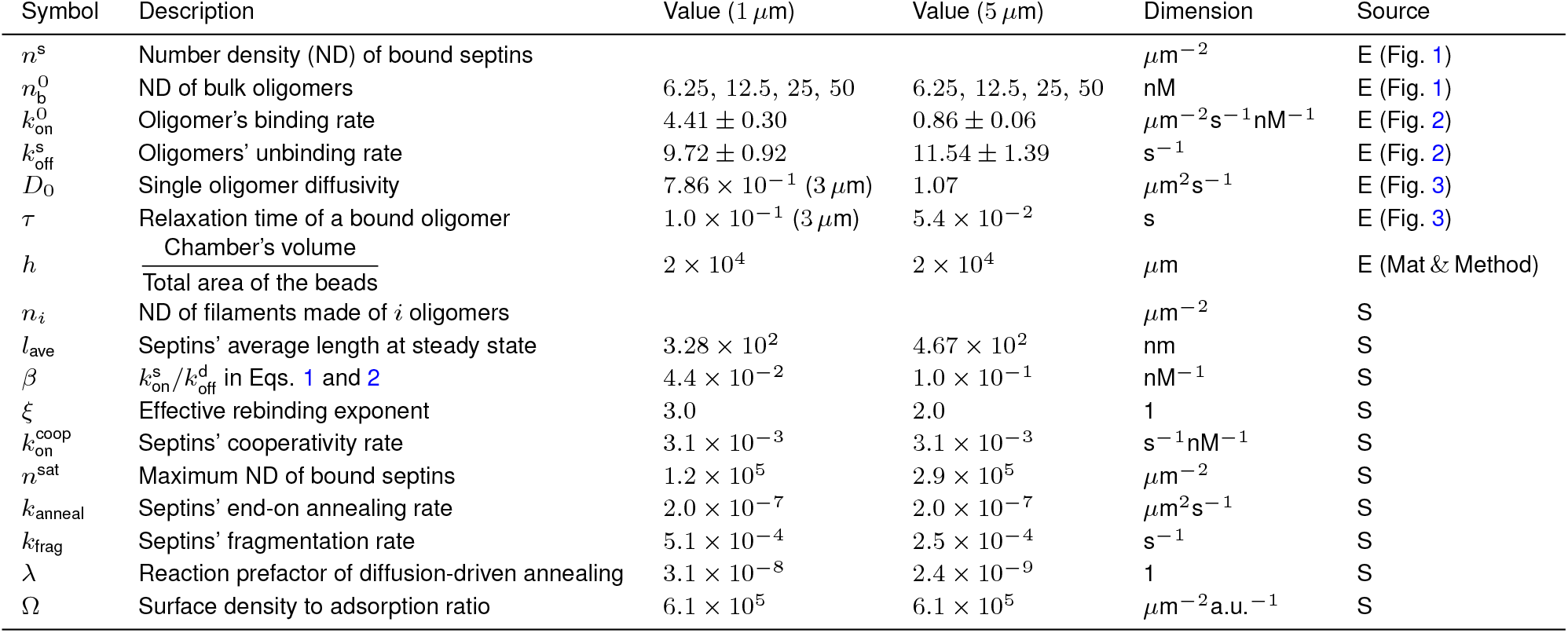
Parameters that appear in the simulations. Parameters labelled as “E” under source column denote variables that are known experimentally and those labelled as “S” (simulation) are determined through least-square optimization of the predictions against experiments.

The typical choice for solving the direct problem is to integrate the overdamped Langevin equation to compute the displacements of septin on the membrane using Eq. 4 in combination with Eq. 6 to describe the kinetic processes. The main challenge in using this method is that the timescale for reaching steady-state adsorption (minutes) is significantly larger than a single oligomer dwell time (≈ 0.1 s), during which the oligomers are undergoing sub-diffusive displacements. As a result, an exceedingly large number of time steps are needed to reach steady-state and resolve the sub-diffusive dynamics of the septins, making this method computationally prohibitive.

As an alternative, we consider the two limits for the annealing process(1) reaction-limited, where the annealing kinetics are controlled by the slower reaction rate when the septins meet; and (2) diffusion-limited reaction, where the annealing kinetics are solely determined by the time it takes the septins to meet. In the reaction-limited case, the annealing reaction of bound septin filaments of length *i* and *j* oligomers is simply given by *dn_i+j_/*dt* = *k*_anneal n_i_n_j__*, where *n_i_* denotes surface density of a filament of length *i* oligomers, and *k*_anneal_ is only a function of bead’s curvature and independent of bulk concentration, filaments’ lengths and surface densities. We use Smoluchowski and Noyes theories (32) in 2D (surface reactions) to model the diffusion-limited reaction rate between two components *i* and *j*. These theories, however, are developed for particles with pure diffusive motion, while the particles in our system are sub-diffusive. The theories can, in principle, be extended to harmonically trapped particles that obey Eq. 4. However, the final expressions are in terms of hypergeometric functions, which are not in closed form and require numerical integration (33). Instead, for simplicity, we use the predicted reaction rate for purely diffusing particles, which presents the maximum reaction rate for our sub-diffusive particles. Following Noyes general theory, we assume only a fraction of encounters, *λ* < 1, result in successful reaction. Including parameter *λ* allows for a simple way of modeling the reaction rate between *i* and *j* components after they meet through diffusive motion: *λ* =1 and *λ* ≪ 1 denote extremely fast and extremely slow reaction rates. The effective reaction rate for this system is approximately

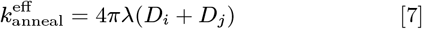

where *D_i_* and *D_j_* are the known diffusion rates of components *i* and *j*, which leaves *λ* as the only fitting parameter in the diffusion-limited reaction model.

Substituting Eq. 7 into the annealing equations, and then the equations in Table 1 into Eqs.6 yields a set of equations that can now be numerically integrated to compute the density of septins of different lengths, *n_i_*, as a function of time. As a result, we can compute the probability density distribution of septins of different length and, thus, the total surface density of bound septins (marked as adsorption here) vs time for a given set of parameters. We can then search for a combination of parameters that gives the least error between the experiments and simulation results given a fitness function.We use the *nonlinlsq* function in Matlab for our optimization process.

## Modeling and Simulation Results

Our kinetic model involves several processes and parameters to be determined through comparison with experiments. The principle of parsimony requires the models to only contain parameters/processes that are necessary for predicting the observations and nothing more. Over-fitting occurs when the model does not meet this standard. Below we provide an argument as to answer why our model is not over-fitting the data. A more comprehensive analysis of the importance of each kinetic process and the sensitivity analysis of the predicted parameters are provided in the later sections and in sections H and I (Figs. S.9-S.21) of supplementary materials.

As shown schematically in Fig. 1D, the time-dependent adsorption curve in each bulk concentration and curvature is primarily characterized by three quantities: the lag time, the slope at the inflection point, and the steady-state adsorption. These variables are experimentally known for each curve; thus, we have 8 × 3 = 24 kinetic data for mono-dispersed assays and 2 × 3 = 6 kinetic parameters for bi-dispersed ones, making the total of 30 meaningful kinetic data. The modeling parameters that need to be determined through least-square fitting are *k*_anneal_ × 2 for reaction limited model (or *λ* × 2 for diffusion-limited model), *n*^sat^ × 2, 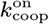, *β* × 2, *k*_frag_ × 2, *ξ* × 2 and *Ω* × 2, making the total of 13 unknown parameters. The symbol “ ×2” denotes that different values are assumed for two different curvatures except for Ω where “×2” denotes mono-dispersed and bi-dispersed case. Note that the number of unknowns (13) is considerably smaller than the number of the experimental kinetic data-points. We determine these 13 unknown through least-square fitting to experimental results. Instead of only using the the kinetic data-points, we use all the measurements of time-dependent adsorption as our experimental data and seek to minimize the error between the predicted and measured adsorption at the same time-steps. Furthermore, the optimization is considered successful only when the predicted average length is in the range 32 nm < *l*_ave_ < 1000 nm and only when the average length is reduced with increasing curvature.

The top row of Fig. 5A and 5B compare the predictions of the model against the measured time-dependent adsorption on mono-dispersed beads of diameter 1 *μ*m and 5 *μ*m, each at different bulk concentrations, where we have used reaction-limited model for the annealing process. The top row of Fig. 5C shows the same comparison for bi-dispersed assays at *n*_b_ = 25 nM bulk concentration. The predictions are displayed with thick solid lines of the same color as their associated experimental data. The computed values of the model are presented in Table 2. Our model gives a reasonable prediction of all three aspects of the assembly process: initiation, growth and saturation in both assays.

**Fig. 5.**
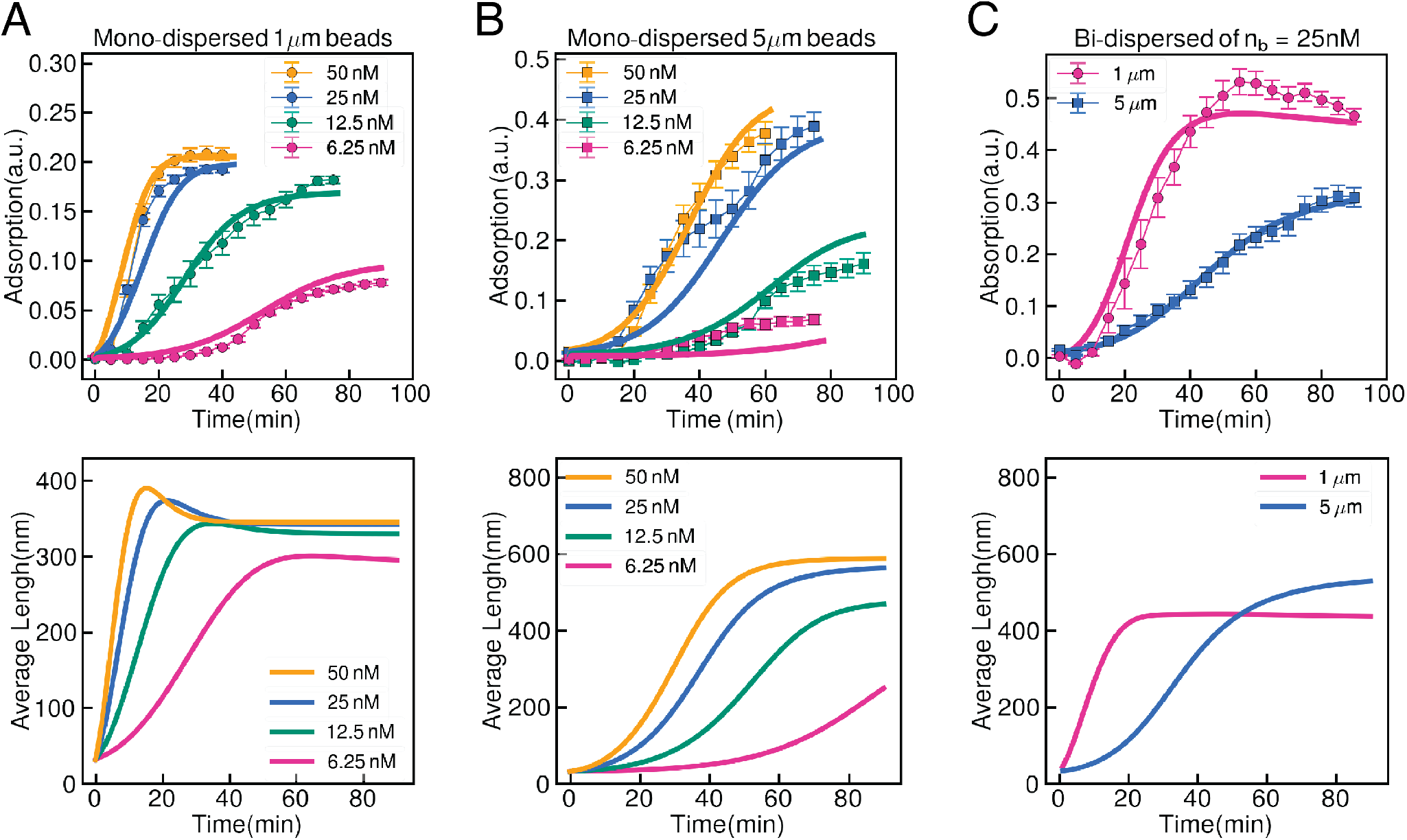
Modeling predictions are in good agreement with time-dependent adsorption experimental data. Top row: Experimental data repeated from Fig. 1 (A-B) Septin adsorption over time in mono-dispersed assays on 1*μ*m (A) and 5*μ*m (B) beads. The filled symbols denote experimental data and solid lines represent the predictions of our kinetic model. Error bars correspond to standard error of th experimental data. (C) Septin adsorption on 1*μ*m (circle symbols) and 5*μ*m (square symbols) beads over time in bi-dispersed assays at 25 nM bulk septin concentration. Bottom row: the variations of the average length of septins vs time, corresponding to the same conditions as the top row.

We also use the diffusion-limited approximation to model the annealing process. Recall that in diffusion-limited model we include the reaction rate by multiplying the effective reaction rate from diffusion by a constant *λ*. The computed value through least-square fitting is 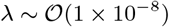, which suggest that effective annealing rate is determined by the rate by which two bound septins merge after they meet, and not by their first passage time that is determined by their relative diffusion. In Fig.S8 we compare the predicted values for lag time, inflection slope and steady-state adsorption for each curve against their experimental values. We see a general agreement between the predicted and measured values. The predicted values for the mentioned parameters all behave monotonically with bulk concentration. The experimentally measured lag times for 5 *μ*m assays show abrupt fluctuations with concentration, which may be due to insufficient sample size (For the same total area of beads, the number of beads and thus the sample size for 5 *μ*m is about 25 times smaller than 1 *μ*m assays) and experimental noise.

Another output of our simulations is the time-dependent length distribution of septins. In Fig.S6 we show the predicted time-dependent length distribution functions in mono- and bi-dispersed assays at different points in time for a selected number of bulk concentrations and curvatures. We observe that after initial times, for all concentrations and for both mono- and bi-dispersed assays the length distributions resemble a gamma distribution. This is also consistent with the length distribution of bound septins on membrane-coated rod assays shown in Fig.S3 in the supplementary materials. The bottom row in Fig. 5 presents the variations of septins’ average length with time (computed from the length distribution function) corresponding to the same conditions at the top row. In 1 *μ*m mono-dispersed assays the average length shows a local maximum very near the inflection point (maximum slope) in its corresponding time-dependent adsorptions. This can be understood by noting that beyond the inflection point the net binding rates start to decrease due to a combination of approaching the saturation concentration and further depletion of bulk septins. Meanwhile the fragmentation and unbinding rates remain unchanged, leading to a net decrease in septins’ length with time.

Focusing now on the steady-state values of the average length, we see that in both mono-dispersed assays the average length increases weakly with septin bulk concentration, with these variations being more pronounced in 5 *μ*m beads. Furthermore, the average steady-state length of septins on 5 *μ*m beads is larger than 1 *μ*m beads at the same bulk concentration for both mono- and bi-dispersed assays. This is in line with the trend observed for length distribution of septins on rod assays, measured from SEM images (Fig. 4). Recall that the underlying reason for this reduction in length is larger fragmentation rates at higher curvatures (1 *μ*m). Indeed, we observe the predicted rates increase from 2.5 × 10^−4^ s^−1^ at 5 *μ*m to double the value 5.1 × 10^−4^ s^−1^ at 1 *μ*m.

### Competition between beads of different curvatures

We now return to our surprising observation in Fig. 1 that motivated the current study: the density of bound septins on 1 *μ*m beads is larger than 5 *μ*m beads in bi-dispersed assays and septins localize to near 1 *μ*m diameter curvatures in cellular contexts (12), while the opposite is true in mono-dispersed assays when only one curvature is present. As it can be seen in Fig. 5, our model can correctly predict this change in behavior between mono- and bi-dispersed assays. The observed trend can be explained as follows. We know that the adsorption rate on 1 *μ*m beads at early and intermediate times is larger than 5 *μ*m beads, leading to shorter initiation and saturation times. Given that 1 *μ*m beads are competing with the slower kinetics on 5 *μ*m beads in bi-dispersed assays, the majority of bulk septins will bind to 1 *μ*m beads, leading to steady-state adsorption levels that are slightly higher than those in monodispersed assays in comparison. We predict the relatively fast binding events of 1 *μ*m beads in bi-dispersed assays results in considerable depletion of septin oligomers in the bulk. This, in turn, reduces the effective binding flux of bulk septins to 5 *μ*m beads, ultimately leading to lower steady-state adsorption compared to 5 *μ*m mono-dispersed beads and lower than the adsorption of 1 *μ*m beads in the bi-dispersed experiments.

Since this reversal in adsorption in mono- and bi-dispersed assays in our model is driven by the depletion of bulk septins, the model predicts that reducing the extent of depletion levels in bi-dispersed assays would favor maximum adsorption on 5 *μ*m beads, leading to a decrease in the ratio of steady-state adsorptions of 1 *μ*m to 5 *μ*m beads. To test this prediction, we manipulated the degree of bulk depletion by changing the total surface area (total number of beads in the assay) in the reactions. Specifically, we performed the bi-dispersed assay experiment, with 25%, 50%, and 75% percent of the total beads’ surface area of the first experiment. The ratio of steady-state adsorption values of 1 *μ*m to 5 *μ*m beads are shown in Fig. 6. Consistent with our model predictions, the adsorption ratio approaches that of mono-dispersed assays as the depletion extent is reduced at lower surface areas.

**Fig. 6.**
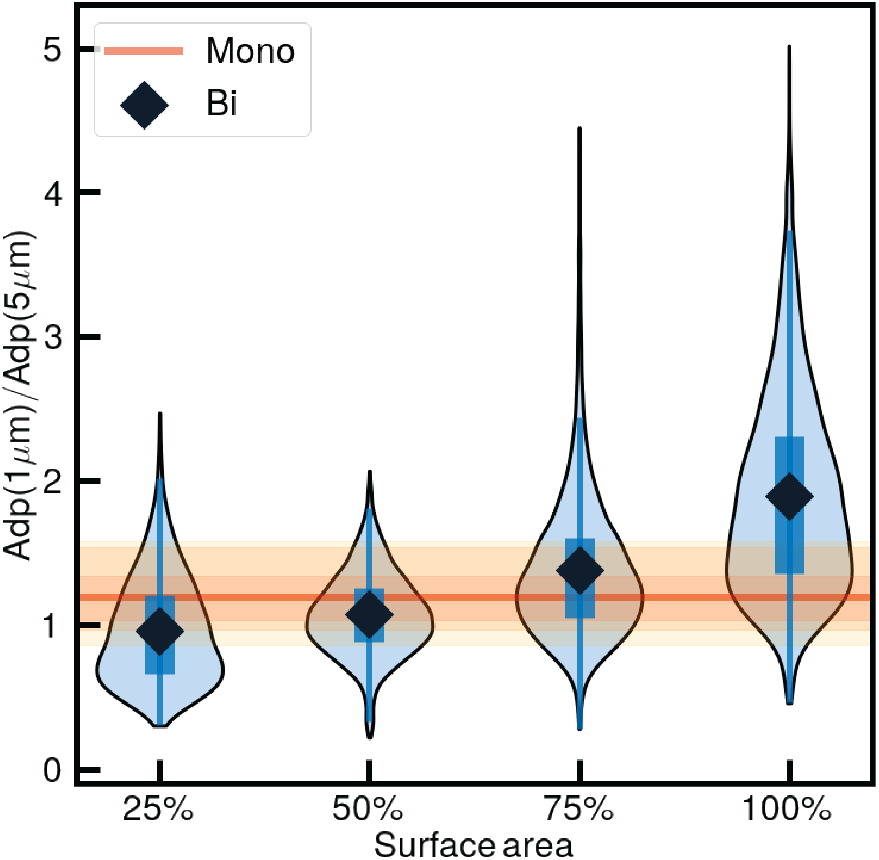
Reducing the degree of septin depletion from the bulk changes septins’ apparent curvature sensitivity. The ratio of steady-state septin adsorption on 1 *μ*m to 5 *μ*m beads in bi-dispersed assays, using different total membrane surface area. Surface area is indicated as a percentage of the standard surface area we used in these assays (5 mm^2^). The bulk septin concentration is 6.25 nM. The blue violin plot shows the distribution of the ratio, and the black diamond indicates the average value. The orange line represents the ratio of adsorption between 1 *μ*m and 5 *μ*m beads at steady state in mono-dispersed assays; the orange belts represent spread in experimental data in the interquartile range (darkest, 25% – 75%), interdecile range (next darkest, 10% – 90%), and full range (lightest, 0% – 100%).

As we mentioned earlier, the measured fluorescent intensity, labeled as adsorption here, is proportional to the surface density of bound septins: *n*^s^ = Ω × Adsorption. Including depletion effects in our model and the change of behavior we observe between mono- and bi-dispersed assays allows us to compute the proportionality constant Ω and, thus, *n*^s^. If we approximate the area occupied by each septin oligomer as a rectangle that is 32 nm in length and 4 nm in width (19), we can also calculate the total area covered by septins. Dividing these numbers by the known total beads’ area gives the area fractions of septins in the experiments. Interestingly, all predicted number of layers are larger than one and in the range 3 – 21, suggesting a multi-layered assembly of septin filaments. In fact, if we assume a maximum packing fraction of 1, these computed surface fractions serve as the lower-bound estimates of the number of septin layers on a membrane surface.

Recent analysis of transmission electron microscopy images of fly and mammalian septins on planar membranes (34) and the results of high-speed atomic microscopy of yeast septins (35) reveal that septins can indeed form multilayered assemblies. However, having 21 layers seems unlikely. Another possibility is that, while the current model assumes all bulk septins have identical binding kinetics, they may consist of sub-populations with different binding kinetics. It is straightforward to modify the model so that, for example, only a fraction *ϵ* of bulk septins can bind and polymerize on the membrane. The adsorption predictions remain unchanged under this modification i.e. we can still predict the depletion effects and the reverse trends observed for absorption in mono- and bi-dispersed assays by simply rescaling the kinetic parameters. The most important difference in predictions is: 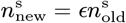. In such a condition, we may observe a single layer of septins, while still observing the novel behaviors that arise from a combination of the depletion of reactive bulk septins and the competition between different curvatures in recruiting these septins from the bulk.

### Septin polymerization is driven by cooperative binding

Next, we asked if including all the elements incorporated in the model, such as cooperativity, annealing, fragmentation, is essential for predicting the correct time-dependent adsorption curves. To test this, we inactivated each of these individual processes in our model and solved for the best agreements between the predictions and experiments. Figures S9-14 in the supplementary materials show the results of time-dependent adsorption and steady-state length distribution under these numerical experiments for both mono-dispersed and bi-dispersed assays. Below we summarize the main observations.

In all instances, aside from end-on annealing, eliminating the processes leads to poor predictions of the time-dependent adsorption curves in mono-dispersed system and even poorer predictions of bi-dispersed adsorptions. Furthermore, excluding fragmentation leads to all bound septins polymerizing to their maximum allowed length in the simulations (the maximum length in our simulations was set to 60 oligomers for 5 *μ*m beads and 40 oligomers for 1 *μ*m beads; changing this maximum length produced almost exactly the same results.). The same trend is observed, when we exclude the cooperative binding. As expected, simulations without depletion completely fail to predict the results of bi-dispersed systems. Setting *β* = 0, which models the competition between septin oligomer binding and defect healing, results in poor predictions of the adsorption curve, particularly the steady-state adsorptions.

Interestingly, we find that our model gives fairly good predictions of time-dependent adsorption after excluding end-on annealing process, and only including the direct and cooperative binding, fragmentation and unbinding processes. Recall that in the absence of end-on annealing bound septins can still polymerize through cooperative binding of bulk septin oligomers to the ends of bound septins. Our modeling results suggest that septin polymerization on curved membranes may be predominantly driven by cooperative binding of subunits from the bulk.

Note that in end-on annealing process filament pairs merge and form longer filaments, without changing the total density of bound septins. Thus, we expect that for a given surface density of septins, the simulations with end-on annealing should produce longer filaments than those without end-on annealing. Figure S15 compare the variations of the average length of septins at steady-state against the steady-state adsorption at different bulk concentration for 1 *μ*m and 5 *μ*m assays, in the presence and absence of end-on annealing. We see that when end-on annealing is included, in both assays, the average length increases roughly linearly with the steady-state adsorption. In contrast, when end-on annealing is excluded, the average length decreases with adsorption in both curvatures. These observations show that the length distribution of bound septins is more sensitive to the details of the assemblies than their net surface density. Experimental measurements of the length distribution at different bulk concentrations would allow us to inspect the relative importance of these assembly processes in more depth and to further constrain the range of modeling parameters.

Finally we perform sensitivity analysis on the parameters computed through optimization. Specifically, we compute the change in the relative errors in the computed values of time lag, inflection slope and steady-state adsorption as the parameters of the model are varied around their optimal values. These results are presented in Figs. S16-21 for both mono-dispersed and bi-dispersed assays. We observe that the relative error are significantly increased for varying the parameters around their optimized value, for at least one of the three variables (lag time, maximum slope and steady-state adsorption). This analysis shows that all parameters except the annealing rate have significant quantitative effect on the time-dependent adsorption, and this combination of computed parameters minimize the error between the experiments and predictions in the wide range of parameter variations explored here.

## Discussion

Micrometer-curvature sensing by nanometer-scale proteins presents a challenge due to the mismatch in length scales. Our previous work (9, 12) had led to a working model where differences in the rates of a single oligomer binding to beads of different curvatures was the basis for septins distinguishing amongst different sizes. This mechanism is also consistent with studies of another micron-scale membrane curvature sensor, SpoVM, that binds the micron-scale forespore membrane in bacteria (10). However, the additional experiments presented here along with the modeling, reveal that while the binding rate is likely a basis for determining initial kinetic differences amongst different curvatures, it is insufficient to support a preference of septins for an optimal curvature. This realization emerged from a simple change in an experimental assay, where we compared curvature-dependent adsorption in regimes where septins must choose between two curvatures or where there is a single curvature present.

Remarkably, depending on available curvature options, different curvature preferences are manifest, suggesting that the curvature sensitivity of septins is not a fixed property of the polymer but instead is sensitive to the process and context of assembly. A series of results presented here reveal that it is the kinetics that determines specific curvature preferences of septins: (1) We showed that single oligomer kinetics alone cannot describe the observed curvature preference of septins, and the assembly of bound septin oligomers must be considered. (2) Our confocal microscopy data shows that septin binding vs time is composed of initiation, growth and saturation periods (Fig. 1D). (3) Using our current and previous experimental results, we developed a kinetic model that describes the time-dependent septin assembly in terms of direct and cooperative binding of septin oligomers and diffusion, end-on annealing, fragmentation and unbinding kinetics of bound septins. (4) We found very good agreements between the kinetic model and experiments, and surprisingly we found that we can do so without the need to include diffusion and end-on annealing kinetics. (5) We observed that the steady-state adsorption on 5 *μ*m beads is larger than 1 *μ*m beads in mono-dispersed assays and this is reversed in bi-dispersed assays of the same bulk concentration. Our model can reproduce this behavior by accounting for competition between fast kinetics on 1 *μ*m beads and slow kinetics on 5 *μ*m beads for a finite pool of septins in the bulk that gets depleted over time. (6) The predicted values of depletion in both assays suggest that septins may be organized into a multi-layered structure on the membrane.

Based on all of these findings and observations we propose the following model for binding and organization of septins on membranes (Fig. 7). The initiation period begins with septin oligomers directly binding to the membrane and through reaction-limited end-on annealing they form longer and more stable filaments. When the densities of bulk and membranebound septins (or cooperative binding rates) are sufficiently large and the unbinding rates are sufficiently small, the bulk septins will bind to the membrane through interactions with the bound filaments giving rise to a superlinear increase in adsorption in growth state. Our predictions in mono- and bi-dispersed assays suggest that this cooperative binding controls the polymerization process.

**Fig. 7.**
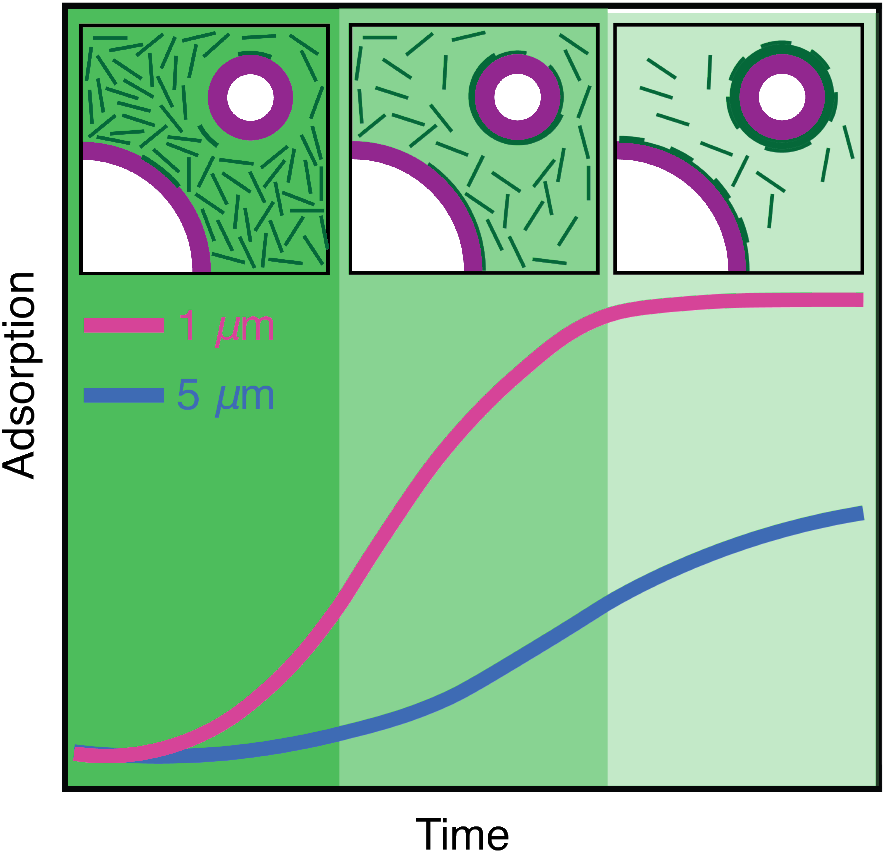
A schematic model of the kinetic basis of septin curvature sensitivity in competition case. Septin adsorption on 1 *μ*m (pink) and 5 *μ*m (blue) beads is plotted over time. Insets illustrate how septins are depleted from the bulk solution primarily by 1 *μ*m beads, leading to faster initiation and growth phases on 1 *μ*m, and ultimately larger adsorptions. Because 5 *μ*m beads recruit from a more depleted bulk, compared to mono-dispersed assays, their growth rate and final adsorption are reduced. Shades of green represent septin bulk concentration and purple represents beads coated with membrane.

In sum, these data point to a key polymer-polymer interaction that builds hiarchial structures of septins as an essential step of curvature-dependent assembly on the micron scale. Importantly, it is possible that control of different binding equilibrium in cells is regulated by other proteins or modifications to the septins that could further tune the kinetics and ultimately the curvature preference of septins. Although in many contexts septins associate with micron-scale membrane curvature, there are wide ranges of curvatures where they bind within this scale that could be dependent on both the binding affinities set by local membrane composition and/or the presence of multiple different curvatures competing for a common pool of limiting septins. This raises the possibility that control of curvature sensing could also emerge through very general mechanisms that change bulk protein concentrations through regulation of transcription/translation/degradation.

Specificity in many biological processes often invokes compatible shapes and electrostatics that govern the thermodynamics of interactions based on structural features. However, there are examples in biology where kinetics gives rise to or supports specificity in a process. Decades ago the concept of kinetic proofreading was proposed for enzyme kinetics and has subsequently been applied to a variety of processes but in essence emerges from varying dissociation rates of nonproductive or inaccurate reaction products (36, 37). Recently the kinetics of association has been invoked for determining sequence specificity in transcription factor binding to specific DNA sequences (38). We find curvature sensitivity in septins to be dependent on multi-step kinetics and by using kinetics rather than the chemomechanical and geometric properties of septin oligomers and membrane, the cell actually can employ these proteins to bind to wider range of possible curvatures. Thus, in this way kinetic-control can provide differential sensitivities within the same molecule, depending on the local geometries present and the abundance of the molecules.

## Materials and Methods

### Saccharomyces cerevisiae septin complex purification

BL21 (DE3) E.coli were co-transformed with plasmids (see below) encoding the budding yeast septins, Cdc11, Cdc12, Cdc3, and Cdc10 and selected for using ampicillin and chloramphenicol. Selected cells were grown in 1 L of terrific broth at 37 °C to an optical density at 600 nm (OD600) between 0.6 – 0.8 at 37 °C. Cells were induced with 1 mM of isopropyl *β*-D-1-thiogalactopyranoside and shifted to 24 °C for 24 h. Cells were harvested by spinning at 10 000 RCF for 20 min. Cell pellets were resuspended in lysis buffer (1 M KCl, 50 mM Hepes pH 7.4, 1 mM MgCl_2_, 10% glycerol, 1% Tween-20, a 1 x protease inhibitor tablet, 20 mM imidazole, and 1mg mL^−1^ lysozyme) and lysed using a probe sonicator. The lysate was clarified by centrifugation using an SS-34 rotor at 20 000 RPM for 30 min. Clarified lysate was filtered (pore size: 0.44 *μ*m) and placed over a column packed with cobalt resin (2 mL resin per liter of culture). The lysate-resin mixture was incubated at 4 °C with gentle mixing for 1 h. The column was washed multiple times with wash buffer (1 M KCl, 50 mM Hepes pH 7.4, 20 mM imidazole). Septins were eluted off the column using 300 mM KCl, 50 mM Hepes pH 7.4, 500 mM imidazole and then dialyzed into 300mM KCl, 50mM Hepes pH 7.4, 1 mM *β*-mercaptoethanol for 24 h. At the initiation of dialysis, 60 *μ*g of Tobacco etch virus protease was added to facilitate the cleavage of the 6x-histadine tag on Cdc12. Septins were then concentrated by centrifugation between 1 and 3*μ*M and passed over a second cobalt column and collected. Purity was assessed using sodium dodecyl sulfate polyacrylamide gel electrophoresis, and protein concentration was measured using a Bradford assay.

### Preparation of small unilamellar vesicles

1,2-dioleoyl-sn-glycero-3-phosphocholine (DOPC), L-*α*-phosphatidylinositol (Liver, Bovine) sodium salt (PI), and L-*α*-phosphatidylethanolamine-N-(lissamine rhodamine B sulfonyl) (ammonium salt) (egg-transphosphatidylated, chicken) (Rh-PE) (all Avanti Polar Lipids) were mixed in chloroform at mol% of 75%, 25%, 0.5%, respectively. Lipid films were generated by evaporating solvent using a stream of argon gas. Residual chloroform was evaporated by taking the lipid film and placing it under a vacuum for at least 3 h. Lipid films were hydrated in 300 mM KCl, 50 mM Hepes pH 7.4, 1 mM MgCl_2_ (at 5 mM) over the course of 30 min at 37 °C with gentle agitation every 5 min. Hydrated films were sonicated (using a water bath sonicator) until the solution became transparent.

### Formation of solid supported lipid bilayers on silica microspheres

Supported lipid bilayers were prepared as previously described (12) Briefly, 50 nmol of small unilamellar vesicles (75% DOPC, 25% PI, 0.5% Rh-PE) were mixed with 440 mm^2^ of silica microsphere for 1 h at room temperature with constant rocking. Lipid-coated beads were pelleted at their minimal sedimentation velocity and washed several times using 33mM KCl and 50mM Hepes pH 7.4.

### Monitoring adsorption of septins onto supported lipid bilayers

Measuring septin adsorption onto lipid-coated silica microspheres was performed as previously described (9, 12). In short, 5 mm^2^ of supported lipid bilayers were mixed with various concentrations of fluorescent septins (final buffer composition of 100 mM KCl, 50 mM Hepes pH 7.4, 0.1 mg mL^−1^ fatty acid free bovine serum albumin, 0.1% methyl cellulose, 1 mM *β*-mercaptoethanol) in a custom reaction chamber glued onto a polyethylene glycol-passivated coverslip. Images were acquired over time using a custom built spinning disc (Yokogowa) confocal microscope equipped with a Ti-82 Nikon stage, 100 x Plan Apo 1.49 NA oil objective, and a Zyla sCMOS (Andor) camera. Analysis of septin adsorption over time was carried out in Imaris 8.1.2 (Bitplane, AG). Raw images were background subtracted in both channels using the software’s Gaussian filter for determination of background (filter width: 33.1 *μ*m). Regions of interest (corresponding to supported lipid bilayers) were defined using the lipid channel to generate a surface rendering. The raw values for intensity sums for both channels over time were exported into Microsoft Excel. Adsorption is defined as the ratio of the intensity sum of the septin surface divided by the intensity sum of the lipid surface. The mean and standard error of the mean of the adsorption is calculated and plotted over time in Python using Matplotlib.

### Kinetics and diffusion of single septin complexes on supported lipid bilayers

Two polyethylene glycol-coated coverslips were sandwiched together using double sided adhesive tape (Nitto product: 5015ELE) to make 20 *μ*L flow chambers. Non-polymerizable septins and solid supported lipid bilayers were added to the chamber and imaged using a custom built total internal reflection fluorescence microscope (Nikon) equipped with a a Ti-82 Nikon stage, 100x Plan Apo 1.49 NA oil objective, and a Prime 95B CMOS (Photometrics) camera. The number and duration of binding events were preformed manually, whereas particle position over time was determined using Trackmate (27). The mean square displacement and its distribution is calculated and plotted over time in Python using Matplotlib.

### Generation and imaging of septin-rod supported lipid bilayer mixture

Preparation of septin-bound lipid-coated rods is described in (9). Septin-rod mixtures were placed onto a polyethylene glycol-coated 12 mm coverslip and fixed using 2.5% glutaraldehyde in 0.05 M sodium cacodylate pH 7.4 for 30 min at room temperature, followed by two washes in 0.05 M sodium cacodylate pH 7.4. The mixture was post-fixed in 0.5% osmium tetroxide for 30min and washed with 0.05M sodium cacodylate pH 7.4. 1% tannic acid solution was added for 15min, washed out, followed by a second fixation with 0.5% osmium tetroxide for 15min. Osmium was washed out using 0.05M sodium cacodylate pH 7.4. Dehydration of samples was performed by adding increasing higher concentrations of ethanol (30%, 50%, 75%, 100%). Dehydrated samples were mixed with transition fluid (hexamethyldisilazane), airdried, and then desiccated until sputter coating. Samples were sputter coated with a 4 nm layer of gold/palladium (60 : 40) alloy before imaging on a Zeiss Supra 25 Field Emission Scanning Electron Microscope.

## ACKNOWLEDGMENTS

The authors would like to thank Rick Baker, Greg Forest, Ronit Freeman, Daphne Klotsa, and Klaus Hahn for their useful discussions. This work was funded by NIH grant R01GM130934 (ASG), NIH training grant T32GM119999 (KSC, BNC); NSF grant CBET-1944156 (EN); Alfred P. Sloan Foundation grant G-2021-14197 (ASG, EN, CE); and HHMI (ASG).

## Notes

### Competing Interest Statement

The authors have declared no competing interest.

