## Supplemental Material for "A kinetic basis for curvature sensing by septins"

PNAS
www.pnas.org

### Supplementary Information for

#### A kinetic basis for curvature sensing by septins

**Wenzheng Shi, Kevin S. Cannon, Brandy N. Curtis, Christopher Edelmaier, Amy S. Gladfelter and Ehssan Nazockdast**

**Ehssan Nazockdast.**
****

##### This PDF file includes:

Supplementary text
SI References

#### Supporting Information Text

**A. Modeling the sub-diffusive motion of septins on the membrane.** We used Kelvin-Voigt (a parallel spring and dashpot) model as shown in Fig. S.1A to describe the interactions of septins with membranes. The viscous (dashpot) element models the viscous drag induced by motion of septins on the membrane, and the spring element models the attractive interactions between septins' amphipathic domains and the hydrophobic and hydrophilic portions of the lipid bilayer, creating a restorative force that keeps the setpin at a preferred location $\mathbf{x}$. The dynamics of bound septins in the presence of thermal forces is described by the following overdamped Langevin equation for a 2D surface:

$$\gamma \frac{\mathrm{d}\mathbf{x}}{\mathrm{d}t} = \mathbf{F}^{\mathrm{Br}} - K_{\mathrm{sp}}\mathbf{x}, \tag{S.1}$$

where $\mathbf{F}^{\mathrm{Br}}$ is the fluctuating thermal force, $\gamma$ is the drag coefficient of the septin moving on the membrane, and $K_{\mathrm{sp}}$ is the spring coefficient modeling the attractive interactions between AH domain and the membrane. After imposing fluctuation-dissipation theorem the dynamic equation in Laplace ($s$) space is reduced to:

$$s^2\langle\Delta\tilde{r}^2(s)\rangle = 4k_bT\tilde{R}^{-1}(s), \tag{S.2}$$

where $\sim$ denotes variables in Laplace space, and $\tilde{R}(s)$ is the hydrodynamic resistance (response) function that relates force and velocity in $s-$space: $\tilde{F} = \tilde{R}\tilde{U}$, and $\langle\Delta\tilde{r}^2(s)\rangle$ is the MSD in $s-$space. For Kelvin-Voigt model it is straightforward to show: $\tilde{R}(s) = \gamma + K_{\mathrm{sp}}/s$. Upon substitution for $\tilde{R}$ in Eq. S.2 and inversion to real-space we obtain the following equation for MSD:

$$\mathrm{MSD} = 4D_0\tau(1 - \exp(-t/\tau)), \tag{S.3}$$

where $\tau = \gamma/K_{\mathrm{sp}}$, and $D_0 = k_{\mathrm{B}}T/\gamma$ is the diffusion of a single oligomer without AH-membrane interactions (1).

Eq. S.3 describes the MSD of a single oligomer, but it does not specify how the septin length affects its sub-diffusive motion. This is important, as septins polymerize to significantly longer lengths in the experiments. To extend this model to septins composed of $N$ oligomers, we need to specify the functionality of parameters of the model, namely $K_{\mathrm{sp}}$ and $\gamma$, with $N$. The drag coefficient of filament embedded in a spherical membrane depends on its length, membrane viscosity, viscosity of surrounding bulk fluid, and the sphere radius (2). In our case, the ratio between 2D membrane viscosity to 3D viscosity of the solution, which determines Saffman-Delbruick length-scale, is approximately $\ell_0 = 2 - 3\,\mathrm{mm}$, which is significantly larger than the membrane radius and septin lengths. Thus, the drag is independent of $\ell_0$. Furthermore, the ratio between the bound septins' average length (see predictions in the main text and below) to the membrane radius is $l_{\mathrm{s}}/R \sim \mathcal{O}(10^{-1}) \ll 1$. A simple scaling analysis shows that in this regime the drag of a rod-like particle along its axis is nearly independent of the membrane radius and is primarily determined by the friction forces that are induced by the relative motion of the septin rod on the top mono-layer with respect to bottom mono-layer. In this limit, septin's drag coefficient scales linearly with its length: $\gamma \sim N$ (2). We also assume that the effective spring coefficient scales linearly with the number of AH-membrane interactions: $K_{\mathrm{sp}} \sim N$. Upon substitution Eq. S.3 modifies to

$$\mathrm{MSD} = (4D_0\tau/N)(1 - \exp(-t/\tau)). \tag{S.4}$$

As it can be seen while the relaxation timescale of motion remains unchanged, the long-time MSD scales inversely with length ($N$). This trend can be seen in Fig. S.1B-D.

Furthermore, $K_{\mathrm{sp}}$, the spring coefficient modeling the attractive interactions between AH domain and the membrane, is increased with curvature which reduces $\tau = \gamma/K_{\mathrm{sp}}$ and leads to a more subdiffusive motion on curved membranes given the same filament length; see Fig. S.1E.

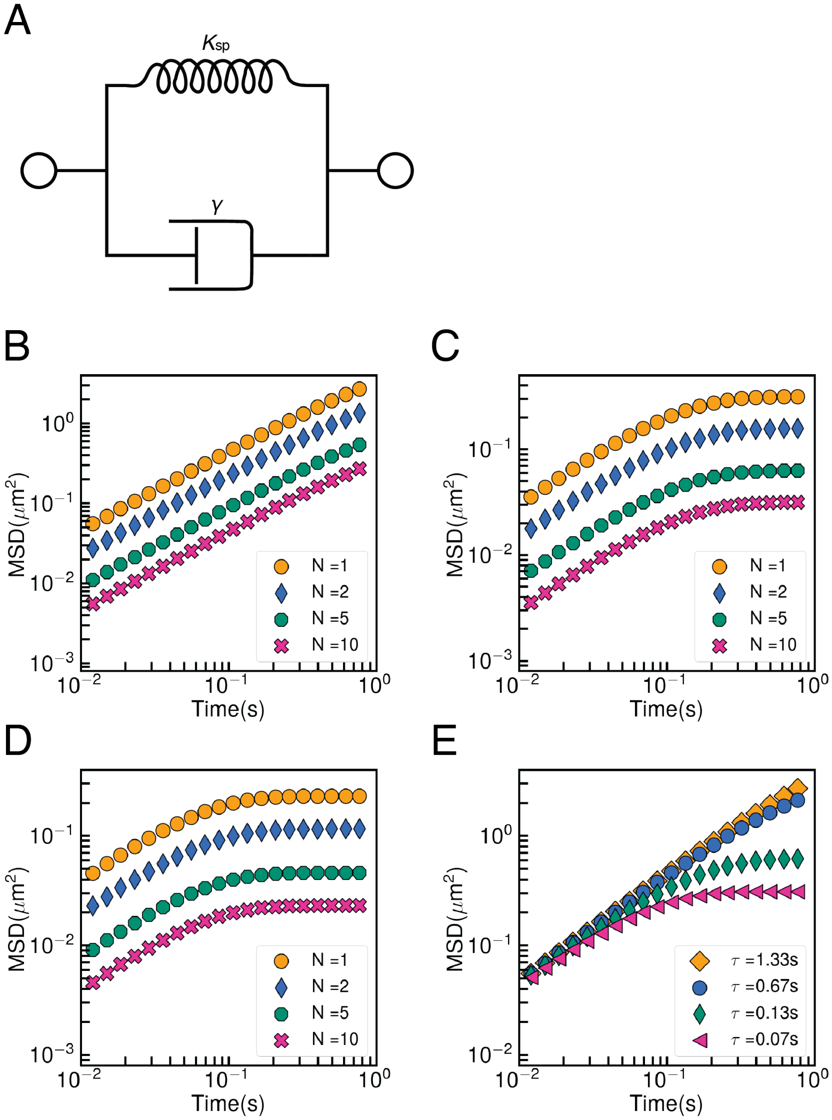


**Fig. S.1.** (A) Kelvin–Voigt model, composed of a parallel spring($K_{\text{sp}}$) and dashpot($\gamma$). (B-D) MSD computed from Kelvin-Void model (Eq. S.4) as a function of time for septin polymers composed of $N$ oligomers on (B) a flat membrane, (C) on membrane curvature $\kappa = 0.67\,\mu\text{m}^{-1}$, and (D) on membrane curvature $\kappa = 0.4\,\mu\text{m}^{-1}$. Different symbols represent different number of oligomers ($N$). The experimental measurements of MSD for single oligomers was used to compute $\gamma$ and $K_{\text{sp}}$ at each curvature. (E) MSD computed from Eq. S.4 as a function of time for septin of single oligomer ($N = 1$) on different curvatures, represented by different $\tau$ here. The experimental measurements of $D_0$ coming from flat membrane was used for all the curves.

**B. Modeling the unbinding rates of septin filaments.** We assume a septin composed of $N$ oligomers can form $N$ independent attachments with the membrane. Each attachment is released with the same rate as a single oligomer: $k^{\rm s}_{\rm off}$. These released attachments can rebind with the rate $k_{\rm rb}$. The dynamics of each attachment is independent of the others. A filament unbinds once all of its attachments are released. The kinetics of the attachment sites can be described by the following system of linear first-order differential equations:

$$\frac{{\rm d}P_1}{{\rm d}t} = -(k^{\rm s}_{\rm off} + (N-1)k_{\rm rb})P_1 + 2k^{\rm s}_{\rm off}P_2, \tag{S.5a}$$

$$\frac{{\rm d}P_i}{{\rm d}t} = (N-i+1)k_{\rm rb}P_{i-1} - (ik^{\rm s}_{\rm off} + (N-i)k_{\rm rb})\,P_i + (i+1)k^{\rm s}_{\rm off}P_{i+1}, \quad (i = 2, 3, \ldots, N-1) \tag{S.5b}$$

$$\frac{{\rm d}P_N}{{\rm d}t} = k_{\rm rb}P_{N-1} - Nk^{\rm s}_{\rm off}P_N, \tag{S.5c}$$

where $P_i$ is the probability of observing $i$ attachments in a filament made of $N$ oligomers. We set our initial conditions such that the entire filament is bound, $P_i(0) = 0$ for $i \neq N$ and $P_N(0) = 1$. Once these equations are solved numerically, the ensemble average lifetime of the septin can be computed as $\langle \tau \rangle_N = k^s_{\rm off} \int_0^\infty tP_1(t)dt$. The effective unbinding rate is the inverse of this dwell time. Our numerical results show that the effective unbinding rate in a wide range of $k_{\rm rb}/k^s_{\rm off} = \mathcal{O}(10^{-1} \sim 10^1)$ is an exponentially decreasing function of the septin's length and can be approximated by

$$k_{\rm off} \approx k^{\rm s}_{\rm off} \exp(-\xi(N-1)), \quad \text{in which} \quad \xi = \frac{3.2k_{\rm rb}/k^s_{\rm off} + 3}{k_{\rm rb}/k^s_{\rm off} + 8.5}. \tag{S.6}$$

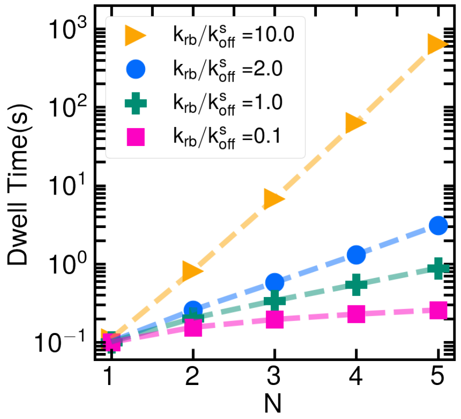


**Fig. S.2.** The dwell time as a function of the number of attachments for different $k_{\rm rb}/k^s_{\rm off}$ rations. Recall that the effective unbinding rate is the inverse of the dwell time.

**C. Post-analysis of the length distribution of septins on membrane-coated rods with different radii.** To study the relationship between septin's length and curvature, we used a previously developed assay of cylindrical rods of radii $R_{\text{rod}} = 46 - 1508\,\text{nm}$, coated with lipids and incubated with septins. Scanning electron microscopy is used to image the bound septins with nanometer resolution. By post-analyzing the SEM images, we measured the length distribution on each rod. Fig. S.3 shows the probability density of the bound septin's length on rods of different radii. Filament lengths range from $32 - 224\,\text{nm}$, which correspond to 1 to 7 oligomers. As a general trend we observe that single oligomers are the most probable length of septins on the rods with smallest radii, while filaments gradually become longer on rods with larger radii. This leads to the average filament length increasing with rod radius. These length distributions follow a gamma distribution for all rod radii, consistent with our predictions.

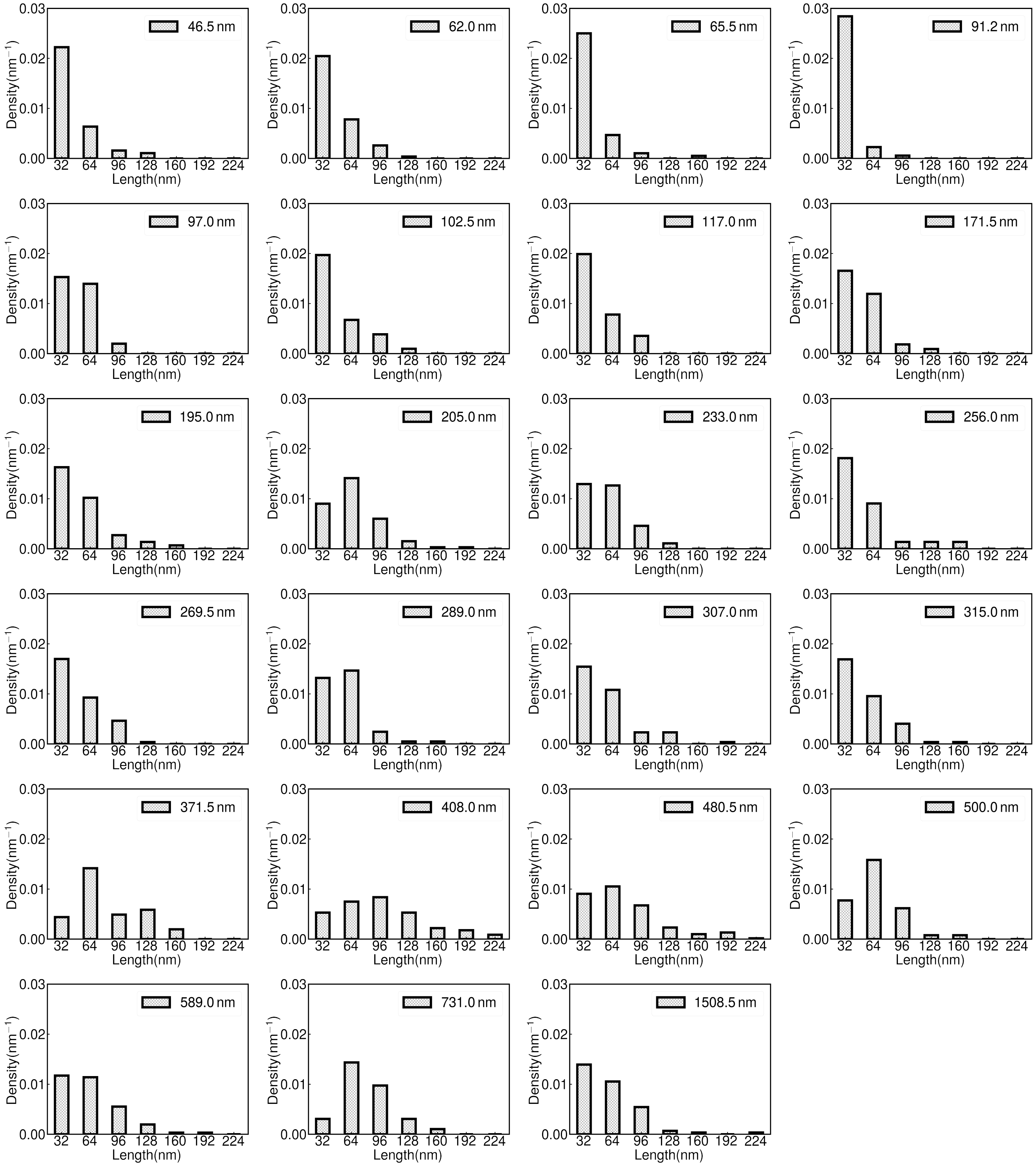


**Fig. S.3.** The distribution of septin filament lengths on rods of different radii. The label represents the radius of the rod.

**D. Measuring septin length vs curvature by using the observed septin alignments on membrane-coated rods.** We used the same rod assay as described in section C to determine how filament length varies its curvature on rods of different radii (Fig S.4). We hypothesized that septins sense the membrane curvature in the direction of their main axis, which is a function of its alignment angle, $\theta$, and rod's radius, $R_{\mathrm{rod}}$: $\kappa = (R_{\mathrm{rod}} \sin\theta)^{-1}$. Using this formula, we computed this curvature for each septin filament seen in the SEM data. We found that oligomers (short filaments) explored a wider range of curvatures, as they can rotate more easily by thermal forces, while longer filaments remained in a narrower curvature range, mostly around $1 - 4\,\mu\mathrm{m}^{-1}$. We also observed that on rods $R_{rod} > 250\,\mathrm{nm}$, the septins predominantly take the maximum allowed curvature $\kappa_{\max} = R_{\mathrm{rod}}^{-1}$, as they do not have access to any larger curvatures. Therefore, when the curvature distribution or average curvature of septins are computed, we exclude the data of $R_{rod} > 250\,\mathrm{nm}$, as those data are biased by the rod radius.

We also observed that the average curvature is independent of the rod's radius, and, thus, independent of any measure of curvature that is solely determined by rod's geometry, such as Gaussian and mean curvatures of the membrane; see Figure S.5A.

Next, we computed the average curvature of septins made of $N = 1 - 4$ oligomers; see Figure S.5B. It appeads that septin's curvature is independent of its length Note that this finding should not be interpreted as septin length is also independent of its curvature.

These observation supports our hypothesis that the curvature that septins sense does not depend on the rod radius nor on the septin filament length, but only depend on the curvature along the septin filament main axis.

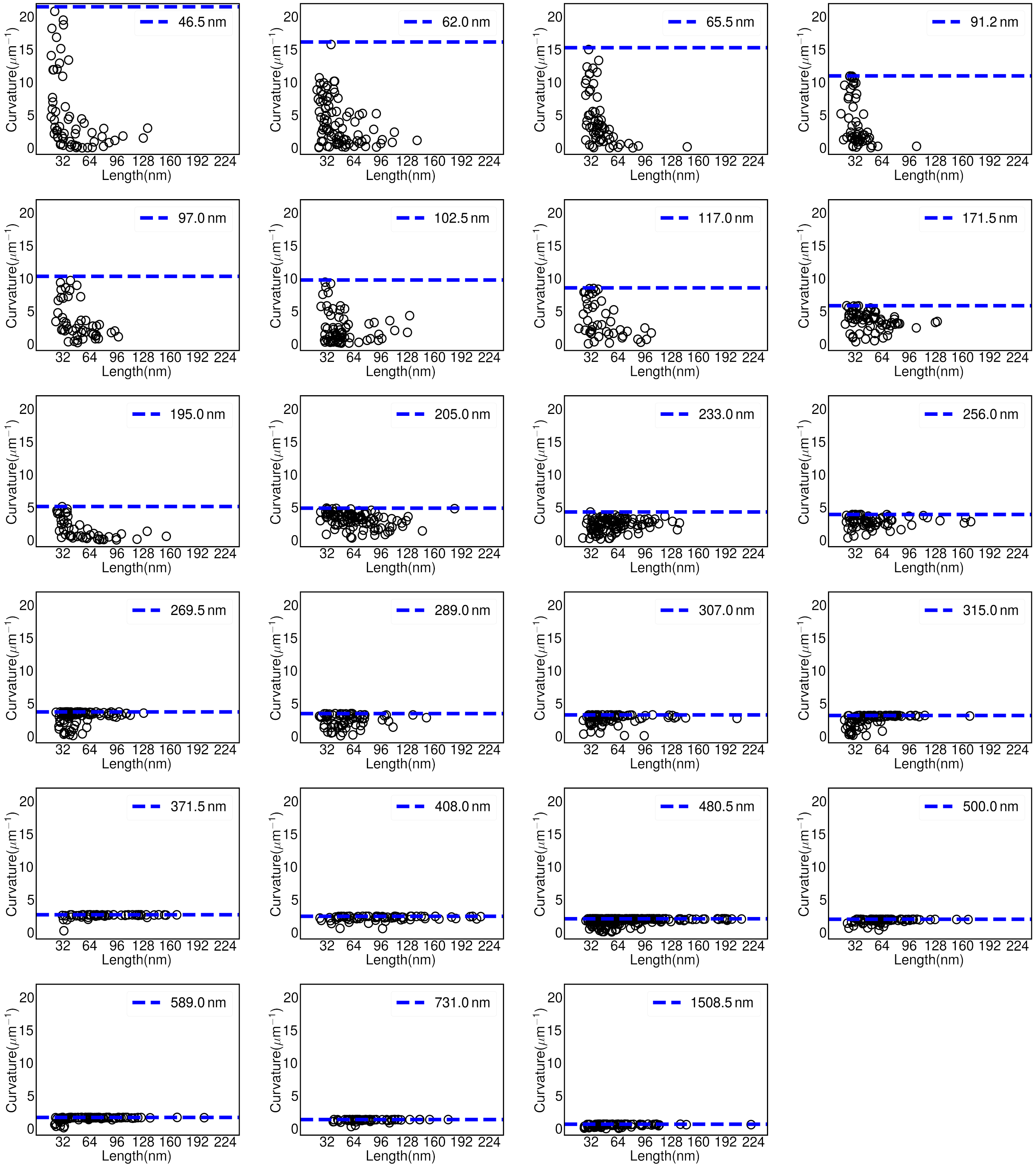


**Fig. S.4.** Observed septin filament curvature as a function of septin filament length on rods of different radii. Each dot represent a septin filament. The blue dashed line is the maximum curvature the septin filament could reach on each rod (inverse of the rod radius).

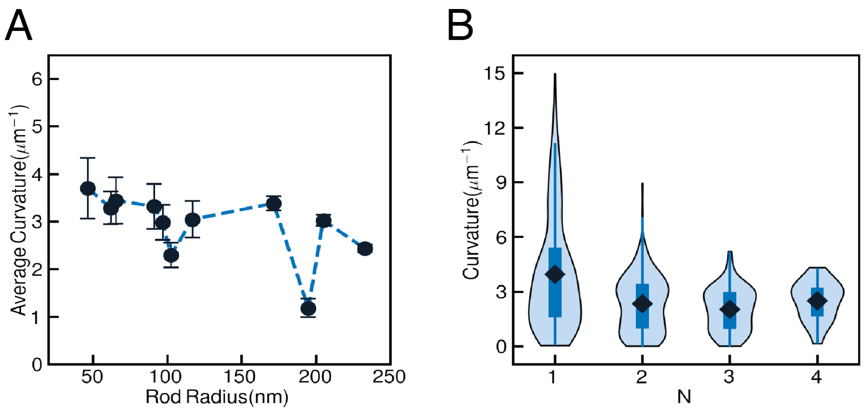


**Fig. S.5.** (A) The average curvature of septins on different rod radii. Dot shows the mean value and error-bar is the standard error. (B) The curvature distribution of septins composed of $N$ oligomers on rod. Diamond symbols highlight the mean.

**E. The predictions of septin filament lengths as a function of time.** Figure S.6 shows the predicted probability densities of septins' length at different instances, for mono- and bi-dispersed systems. Septins bind as oligomers at early times, evident from the strong peak at $N = 1$ (Fig.S.6). As time progresses, septins form longer filaments through a combination of end-on annealing and cooperative binding of bulk septins to the ends of existing bound septins. As a result, the length distribution peak is shifted to larger values of $N$. The probability resembles a gamma-distribution for these intermediate and final times.

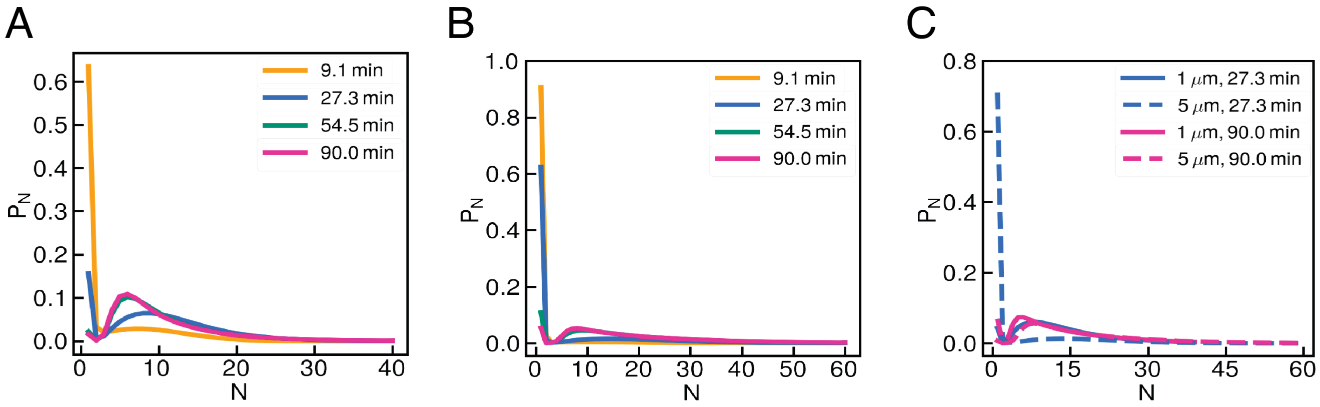


**Fig. S.6. Predicted septin filament lengths as functions of time.** (A) Mono-dispersed system $1\,\mu$m beads of $12.5$ nM at different times. Color in the legend depicts which time point of the distribution. (B) Mono-dispersed system $5\,\mu$m beads of $50$ nM at different times. Color in the legend depicts which time point of the distribution. (C) Bi-dispersed system of $25$ nM. Color and line type in the legend depicts which bead size and which time point was used for the distribution.

**F. Lag time, maximum slope and steady-state values of adsorption in experiments and simulations.** As shown schematically in Fig. 1D in the main text, the time-dependent adsorption plots resembles a sigmoidal curve that can be characterized by three parameters: (1) the initiation or lag time, $T_{\mathrm{lag}}$, (2) the maximum slope, $S_{\mathrm{max}}$, and (3) the steady-state adsorption (saturation plateau), $n_{\mathrm{p}}$ (Fig. S.7). Note that the lag time ($T_{\mathrm{lag}}$) is the intersection of the maximum slope with the time axis. We compute these parameters for each combination of bulk concentration and bead curvature (radius) in simulations and experiments by fitting the following function to the time-dependent adsorption curves:

$$n(t) = \frac{1 - \exp\left(-a(b-c)t\right)}{b^{-1} - c^{-1}\exp\left(-a(b-c)t\right)}, \qquad \text{[S.7]}$$

in which $n(t)$ is the septin adsorption as a function of time, and $a$, $b$, and $c$ are fitting parameters. As shown in Fig. S.7, Eq. S.7 can accurately fit the experimental data. Given the analytical form of the fitting function, we can find a closed form relationships for the lag time, the maximum slope, and the steady-state adsorption in terms of $a$, $b$ and $c$:

$$T_{\mathrm{lag}} = \frac{\ln(-bc^{-1})(b-c) - 2(b+c)}{a(b-c)^2}, \qquad S_{\mathrm{max}} = \left(\frac{\mathrm{d}n}{\mathrm{d}t}\right)_{\mathrm{max}} = \frac{1}{4}a(b-c)^2 \qquad n_{\mathrm{p}} = b. \qquad \text{[S.8]}$$

Fitting Eq. S.7 to simulation results also produces very good agreement. We use the fitting to calculate the lag time, $T_{\mathrm{lag}}$,

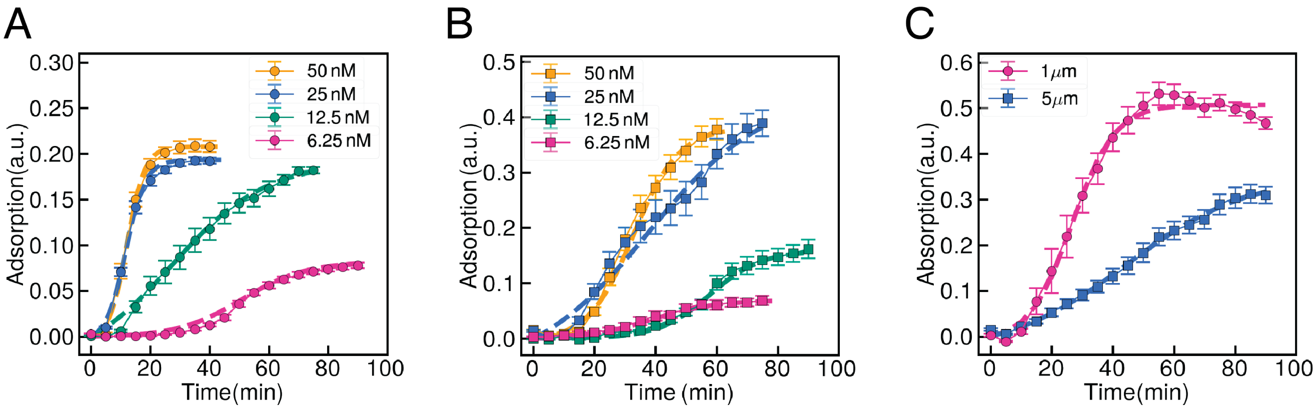


**Fig. S.7. The fitting results and their comparison against the measured time-dependent adsorptions**. The time-dependent adsorption from experiments (symbols with error-bars, where different colors denote different bulk concentrations) against the fitting results using Eq. S.7 (dashed lines of the same color). The left (A) and middle (B) are the results of mono-dispersed $1\ \mu$m and $5\ \mu$m bead assays, and the right (C) shows the results for the bi-dispersed case at a bulk concentration of $25$ nM.

maximum slope, $S_{\mathrm{max}}$ and steady-state adsorption values $n_{\mathrm{p}}$ in the predicted adsorption curves and compare them against their associated experimental values in Fig. S.8. We can see a general agreement between the predicted and experimental values.

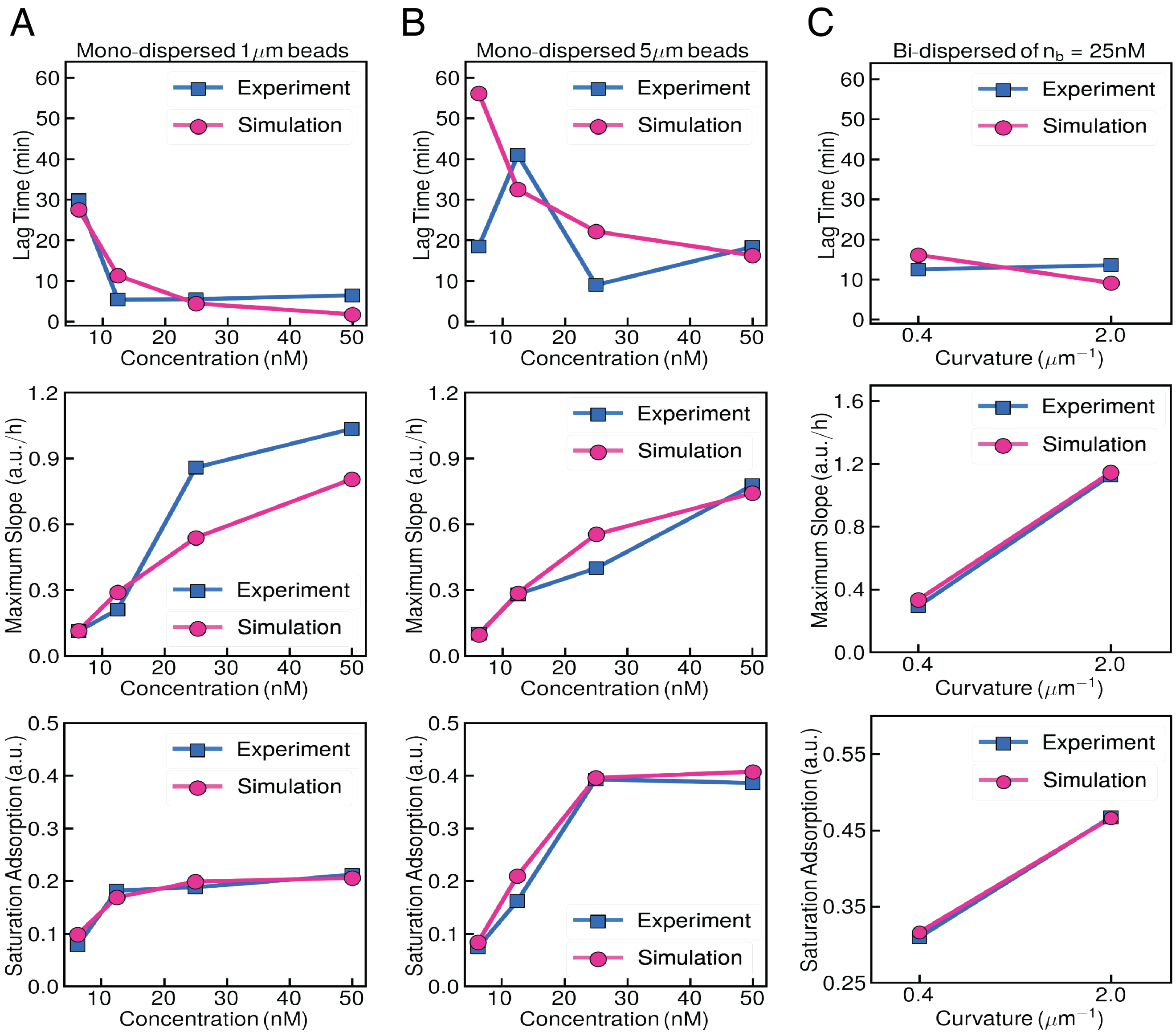


**Fig. S.8. Comparison between the experimental and simulation values for lag time, maximum slope and steady-state adsorption values for time-dependent adsorption**.The left (A), middle (B), and right (C) column represent the mono-dispersed case on $1\ \mu$m, mono-dispersed case on $5\ \mu$m, and bi-dispersed case at a bulk concentration of $25$nM, respectively. The first row is lag time as defined in Eq. S.8. The second row is maximum slope as defined in Eq. S.8. The third row is saturation adsorption at steady state as defined in Eq. S.8. The blue curve and the square mark represent the experiment data while the red curve and the dotted mark represent the simulation.

**G. Assessing the importance of different kinetic processes on the predicted adsorption and length distributions.** Our model incorporates several kinetic processes, including defect formation/healing dynamics, cooperative binding, end-on annealing, fragmentation, and bulk depletion. To examine the importance of each individual process on the overall predictions, we inactivate each process in our model and then solve for the best agreement between the predictions and experiments for mono-dispersed $1\,\mu$m and $5\,\mu$m systems. The parameters computed from this optimization are then used to predict the time-dependent adsorptions and length distributions in mono-dispersed and competitive assays. Our reference set of predictions for the full model are shown in Fig. S.9. The results of these numerical perturbations are provided in Fig. S.10 to S.14. In all figures the left and middle columns compares the predictions of the time-dependent adsorption (top row) against the experimental results and the predictions of the probability density distribution of septins length at the end of simulation time (bottom row) for $1\,\mu$m, for $5\,\mu$m mono-dispersed assays; the figure on the right column show the same results and comparisons in the competitive (bi-dispersed) case. A brief description of the results are provided in the same page and subsection of the relevant figure. Note that the length distribution predictions correspond to the last simulation time-step. The amount of simulation time exactly replicates the time frame in experiments.

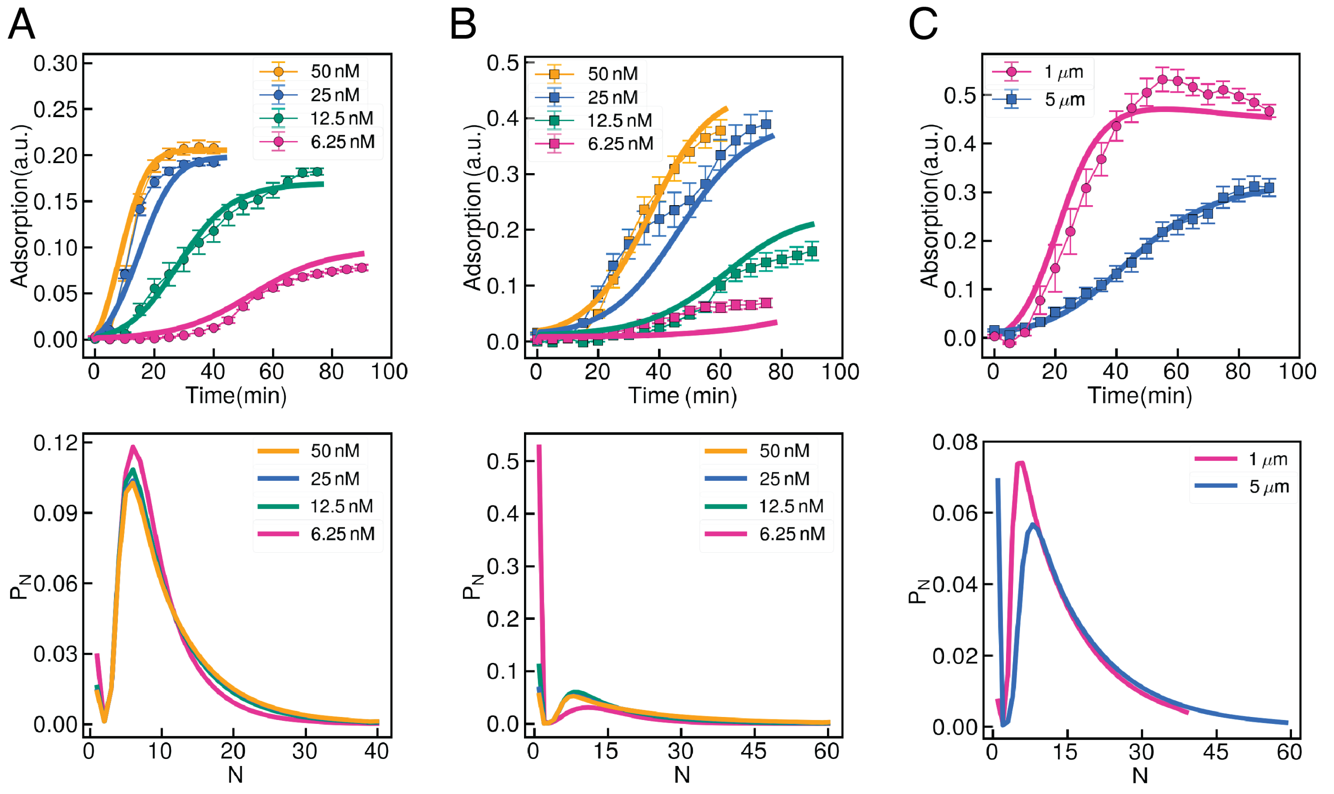


**Fig. S.9. Septin adsorption over time including all the mechanisms.** The top row compares the time-dependent adsorption from experiments (symbols with error-bars, where different colors denote different bulk concentrations) against the simulation results (solid lines of the same color), when all the mechanisms are included. The left (A) and middle (B) columns are the results of mono-dispersed $1\,\mu$m and $5\,\mu$m bead assays, and the right column (C) shows the results of the bi-dispersed case at a bulk concentration of $25$ nM. The bottom rows show the steady-state probability density of septins' length (unit length$^{-1}$) corresponding to the same assays. Error bars in the experimental data correspond to their standard error.

***G.1. Inactivate cooperative binding ($k_{on}^{coop} = 0$).*** Comparing the top rows in Figs. S.9 (all included) and S.10 shows that inactivating cooperativity leads to only slight changes in prediction of the adsorption versus time. In contrast, the predicted steady-state length distributions are very different. In simulations without cooperative binding the most probable length is the maximum allowed length in the simulations, which are respectively 40 and 60 oligomers in $1\,\mu$m and $5\,\mu$m cases. This prediction is in clear contrast with the length distribution of septins in the rod assays. As cooperative binding is inactivated, the adsorption increase requires a much larger end-on annealing rate to form longer filaments which are more stable on the membrane. We note the total adsorption remains unchanged when two filaments merge and form a longer filament in the end-on annealing process. If the end-on annealing rate is sufficiently large and thus leading to slow unbinding rate, the bound filaments will continue to merge and form more stable and longer filaments without any change in adsorption. Thus, even when the adsorption appears to have reached a steady-state, the filaments can continue to increase in length through end-on annealing.

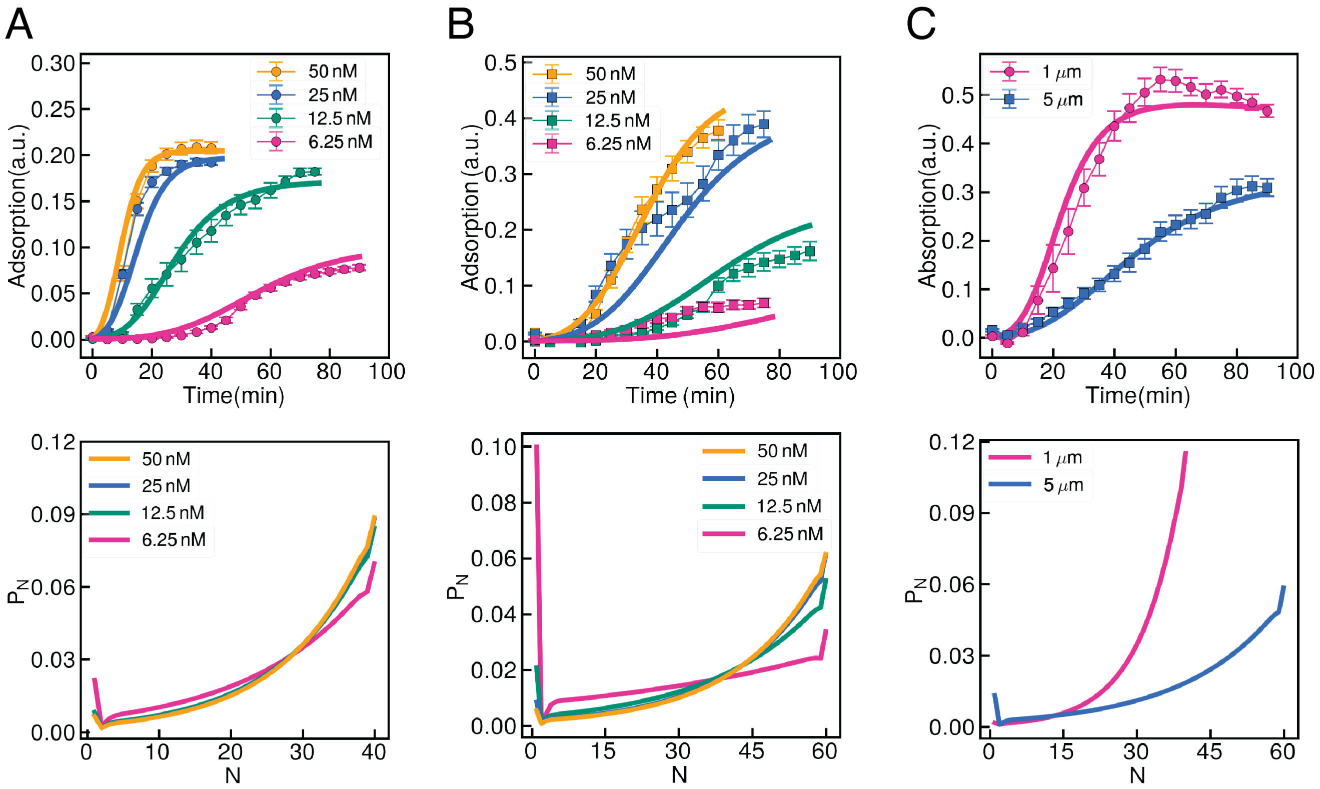


**Fig. S.10. Septin adsorption over time without cooperative binding**. The top row compares the time-dependent adsorption from experiments (symbols with error-bars, where different colors denote different bulk concentrations) against the simulation results (solid lines of the same color), when cooperativity binding is inactivated. The left (A) and middle (B) columns are the results of mono-dispersed $1\,\mu$m and $5\,\mu$m bead assays, and the right column (C) shows the results for the bi-dispersed case at a bulk concentration of $25\,$nM. The bottom rows show the steady-state probability density of septins' length (unit length$^{-1}$) corresponding to the same assays. Error-bars in the experimental data correspond to their standard error.

***G.2. Inactivate filament fragmentation ($k_{frag} = 0$) .*** As it can be seen in Fig. S.11, inactivating fragmentation leads to slightly poorer predictions of time-dependent adsorption compared to the full model (see Fig. S.9). More importantly, inactivating fragmentation leads to unbounded polymerization of all the septins to maximum allowed length in the simulation, which are 40 and 60 oligomer units in $1\,\mu$m and $5\,\mu$m beads. These predictions disagree with the experimental observations of septins' length distribution on membrane-coated rods, discussed both in the main text (see Fig. 4D in the main text) and in the Supplementary materials (see Fig. S3).

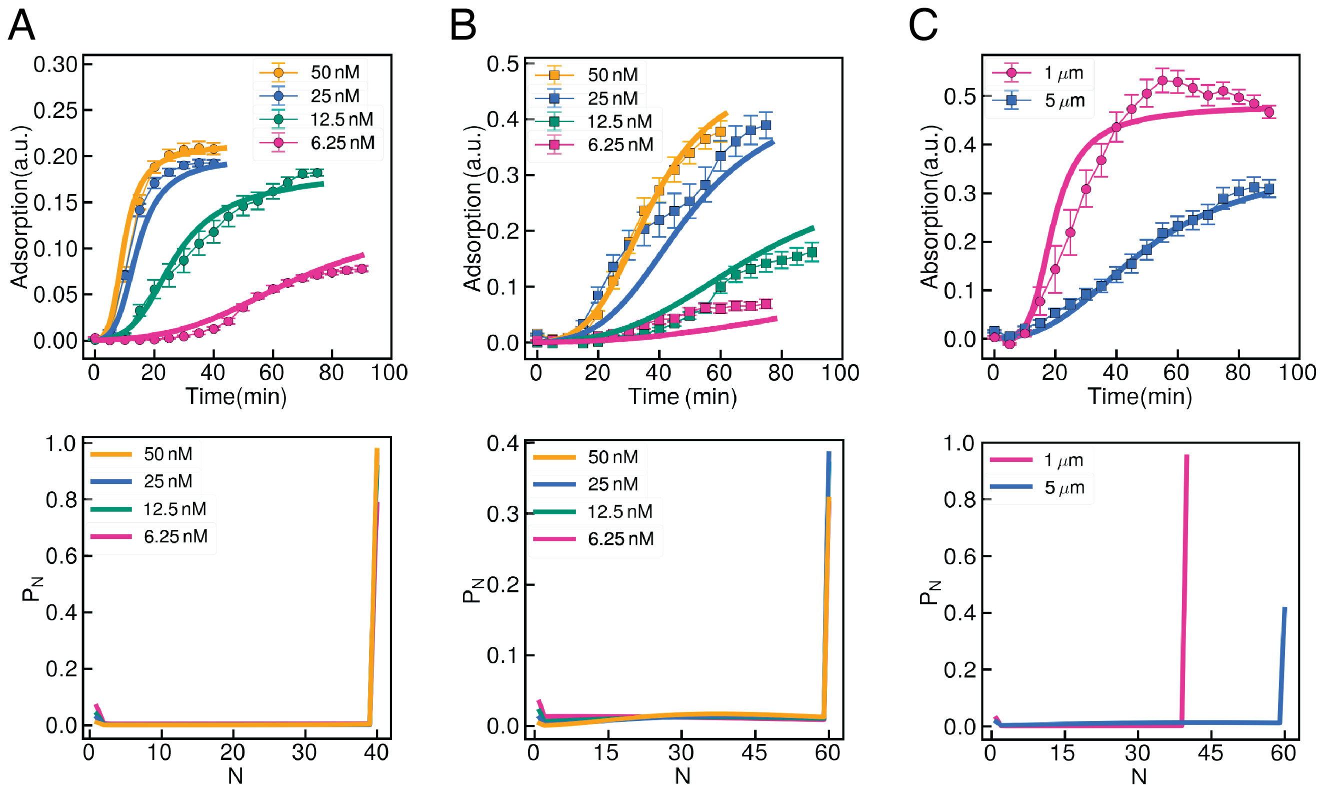


**Fig. S.11. Septin adsorption over time without fragmentation** The top row compares the time-dependent adsorption from experiments (symbols with error-bars, where different colors denote different bulk concentrations) against the simulation results (solid lines of the same color), when fragmentation is inactivated. The left (A) and middle (B) columns are the results of mono-dispersed $1\ \mu$m and $5\ \mu$m bead assays, and the right column (C) shows the results for the bi-dispersed case at a bulk concentration of $25$ nM. The bottom rows show the steady-state probability density of septins' length (unit length$^{-1}$) corresponding to the same assays. Error-bars in the experimental data correspond to their standard error.

***G.3. Inactivate defect dynamics ($\beta = 0$).*** Comparing the predictions against the full model shows that inactivating the competition between defect healing and septin binding dynamics by setting $\beta = 0$ leads to poor predictions of time-dependent adsorption in the mono-dispersed system. As shown in Fig. S.12 A-B, the perturbed model cannot predict the correct initiation time, $T_{\mathrm{lag}}$, especially at large bulk concentrations, i.e. $n_{\mathrm{b}} = 25\,\mathrm{nM}$ and $n_{\mathrm{b}} = 50\,\mathrm{nM}$. Furthermore, as shown in figure S.12C, the simulations with $\beta = 0$ fail to predict the kinetics on $5\,\mu$m bead where kinetics becomes more sensitive to the depletion effect in this scenario.

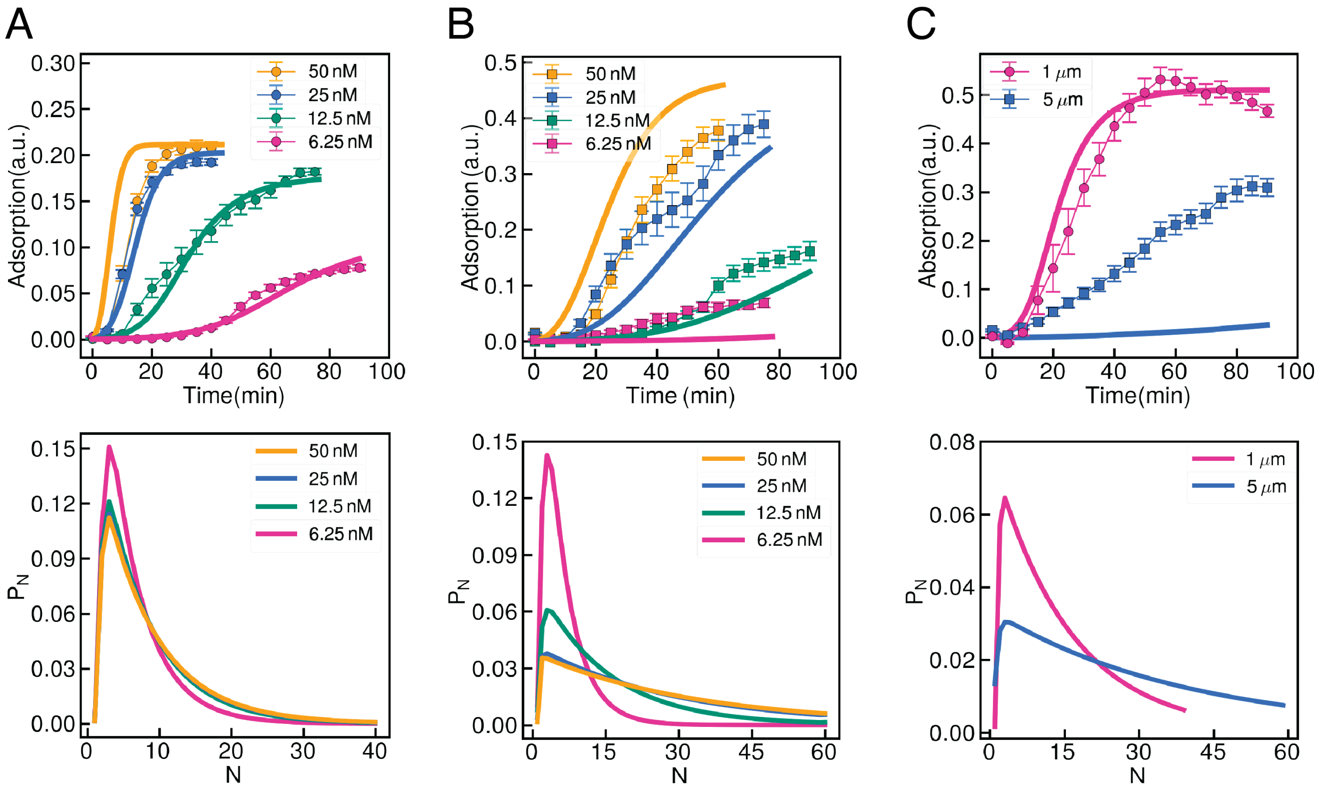


**Fig. S.12. Septin adsorption over time without defect dynamics.** The top row compares the time-dependent adsorption from experiments (symbols with error-bars, where different colors denote different bulk concentrations) against the simulation results (solid lines of the same color), when cooperative binding is inactivated. The left (A) and middle (B) columns are the results of mono-dispersed $1\,\mu$m and $5\,\mu$m bead assays, and the right column (C) shows the results for the bi-dispersed case at a bulk concentration of $25\,$nM. The bottom rows show the density of septins' length (unit length$^{-1}$) at the last simulation time, corresponding to the same assays. Error-bars in the experimental data show their standard error.

***G.4. Inactivate bulk depletion ($A/V = h^{-1} = 0$).*** Comparing the top rows in Figs. S.9 (all included) and S.13 shows that inactivating bulk depletion leads to only moderate changes in prediction of the adsorption versus time in mono-dispersed case (see Fig. S.13 A-B). However, the perturbed model completely fails to predict the bi-dispersed case where the adsorption at steady state is opposite of the experimental result. This is consistent with our argument that the reversal behavior of adsorption amounts in mono- and bi-dispersed assays is caused by the depletion of septins in the bulk. When depletion is inactivated, the kinetics on $1\,\mu$m and $5\,\mu$m become decoupled in the bi-dispersed case; thus, in both mono- and bi-dispersed assays the adsorption is larger on $5\,\mu$m beads.

It is also useful to comment on the predicted peak corresponding to single oligomers in length distribution of $5\,\mu$m mono-dispersed assays with 6.25 nM bulk concentration (the bottom panel of Fig. S.13 B). This behavior is not present in the other conditions. Recall that the length distribution curves correspond to the distributions at the last simulation time-step and may not have reached steady-state. This is certainly true for the case of $5\,\mu$m mono-dispersed assays with 6.25 nM bulk concentration, where the simulations greatly over-predict the initiation time in this bulk concentration. Thus, this length distribution is more suitably categorized as the length distribution at early times. In support of this argument, the predicted length distribution of the full model at early times contains the same features as the one with 6.25 nM bulk concentration i.e. a strong peak at $N = 1$, followed by a weaker peak at intermediate lengths (see Fig. S.6).

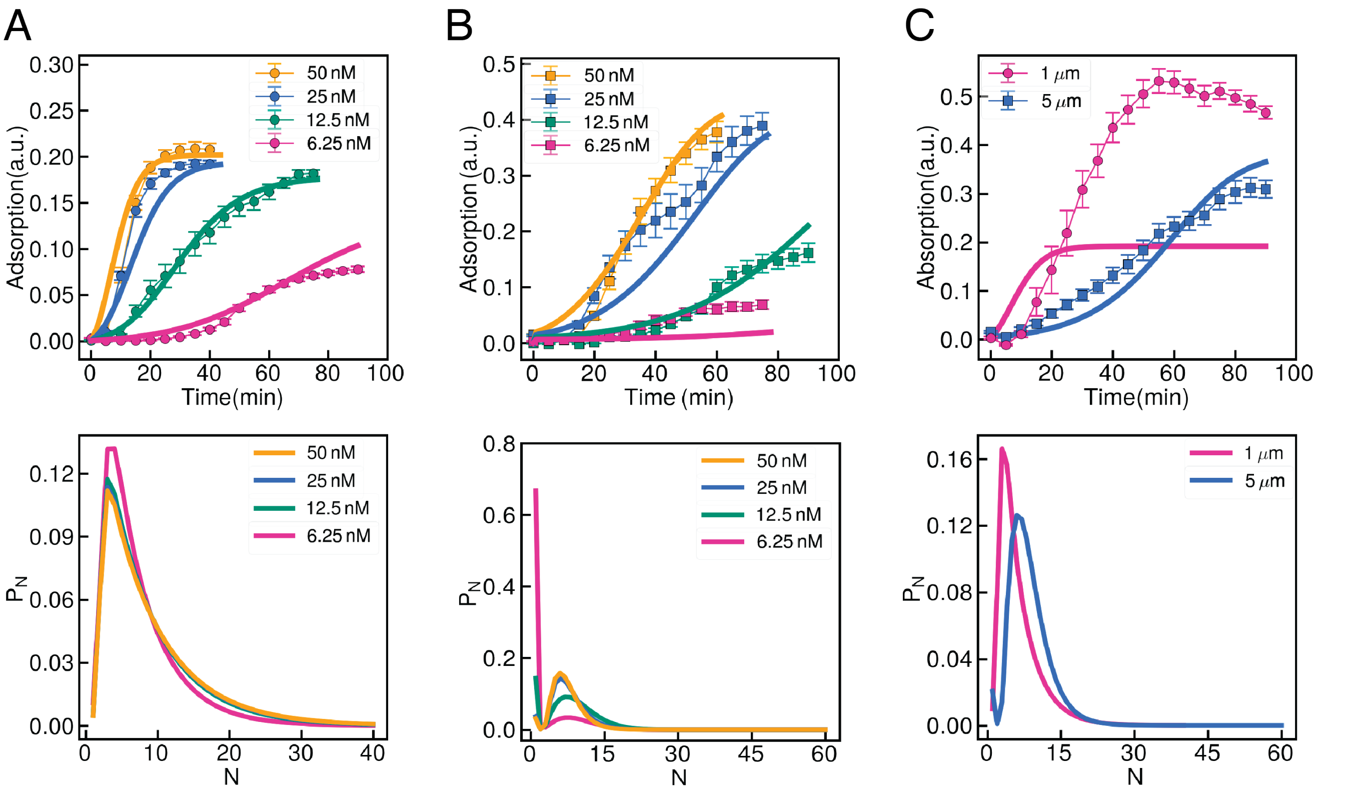


**Fig. S.13. Septin adsorption over time without bulk depletion.** The top row compares the time-dependent adsorption from experiments (symbols with error-bars, where different colors denote different bulk concentrations) against the simulation results (solid lines of the same color), when cooperative binding is inactivated. The left (A) and middle (B) columns are the results of mono-dispersed $1\,\mu$m and $5\,\mu$m bead assays, and the right column (C) shows the results for the bi-dispersed case at a bulk concentration of $25$ nM. The bottom rows show the probability density of septins' length (unit length$^{-1}$) at the last simulation time, corresponding to the same assays. Error-bars in the experimental data show their standard error.

***G.5. Inactivate end-on annealing ($k_{anneal} = 0$).*** End-on annealing is a two-step process involving the (sub)diffusion of bound septins and the reaction between the two ends of bound septins, when they are sufficiently close. We model this two-step process by considering two limits: when the (sub)diffusion timescale is significantly larger than the reaction timescale (diffusion-limited); and when the reaction timescale is the slowest process (reaction-limited). Our full simulation results suggest that the annealing kinetics are determined by the reaction rate (reaction-limited). Hence, to inactive the end-on annealing process we set $k_{\mathrm{anneal}} = 0$. We note that the results remain unchanged if we choose to inactive the diffusion-limited model instead by setting $\lambda = 0$ in that model (see Eq. 7 in the main text), since both variations of the model correspond to no end-on annealing.

A comparison between the predictions of the full model in Fig. S.9 against those in Fig. S.14 shows that the simulation can reproduce the time-dependent adsorption experimental data with roughly the same level of quantitative agreement as the full model. The predicted length distributions are also in the acceptable range, and are in the same order of magnitude as the predictions of the full model. In other words, unlike the other processes, the end-on annealing appears to be dispensable for correctly predicting the experimental observations.

The above observations raise the following question: Does end-on annealing have any experimentally measurable effects on structure and dynamics of septin organization on membranes? Fig. S.15 compares the predictions of the steady-state average length vs steady-state adsorption in the full model for $1\,\mu$m and $5\,\mu$m mono-dispersed assays, against those where annealing is inactivated. Each data-point correspond to different bulk concentrations, bead size and full vs inactivated model. In the full model the average length monotonically increases with the steady-state adsorption, with the slope being larger in $5\,\mu$m beads. In contrast when end-on annealing is excluded, the average length decreases with adsorption in both curvatures, with the slopes being larger in $1\,\mu$m beads.

Bound septins polymerize through either cooperative binding of bulk oligomers to their ends or when two of them merge and form longer filaments in end-on annealing process. While polymerization through annealing keeps the total adsorption unchanged, the polymerization through cooperative binding is always associated with increases in adsorption. This is in line with the observation that length distribution and adsorption are positively correlated (positive slope) in the full model, where polymerization is dominated by cooperative binding, while they show negative correlation when cooperative binding is inactivated. These numerical experiments suggest that measurements of the average length at different bulk concentrations can be used to inspect the relative importance of cooperative binding and end-on annealing and to further constrain the range of modeling parameters.

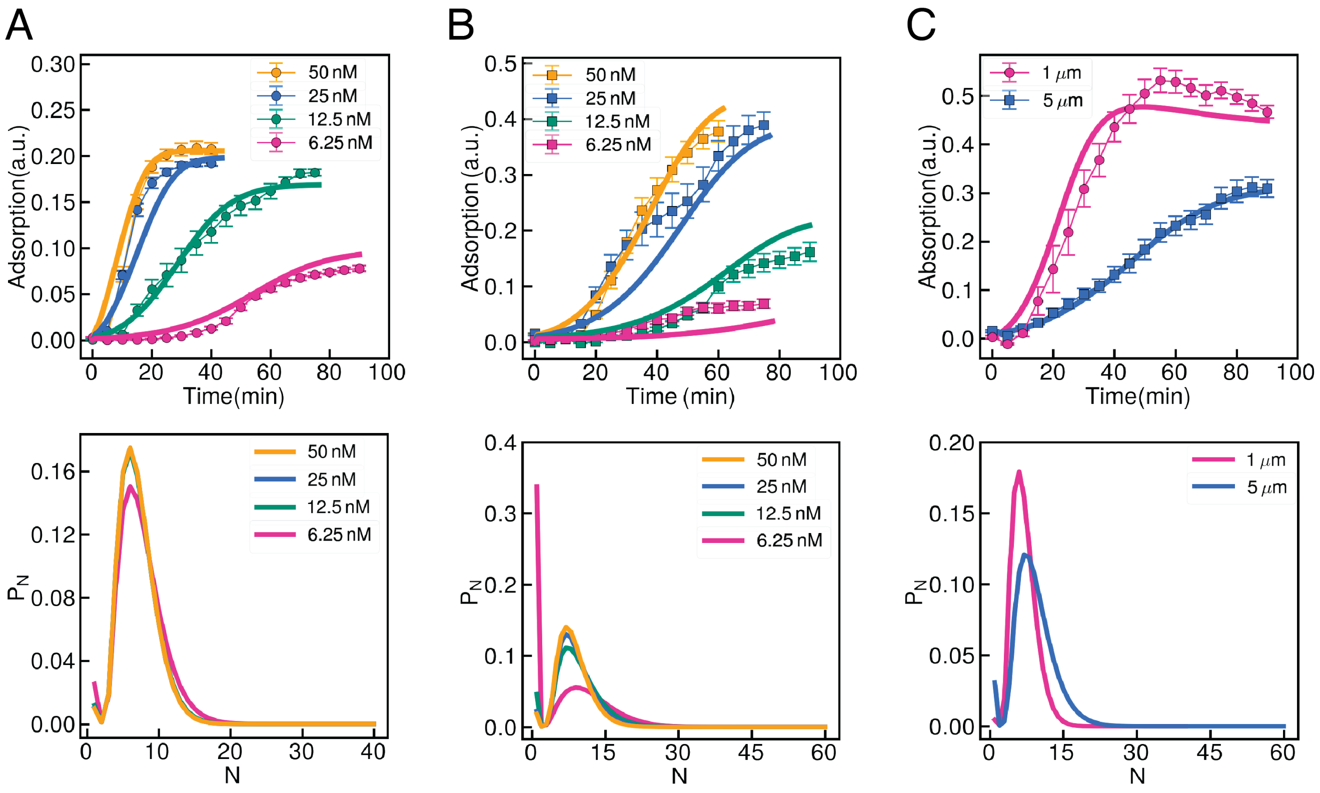


**Fig. S.14. Septin adsorption over time without end-on annealing.** The top row compares the time-dependent adsorption from experiments (symbols with error-bars, where different colors denote different bulk concentrations) against the simulation results (solid lines of the same color), when cooperative binding is inactivated. The left (A) and middle (B) columns are the results of mono-dispersed $1\,\mu$m and $5\,\mu$m bead assays, and the right column (C) shows the results for the bi-dispersed case at a bulk concentration of $25$ nM. The bottom rows show the probability density of septins' length (unit length$^{-1}$) at the last simulation time, corresponding to the same assays. Error-bars in the experimental data show their standard error.

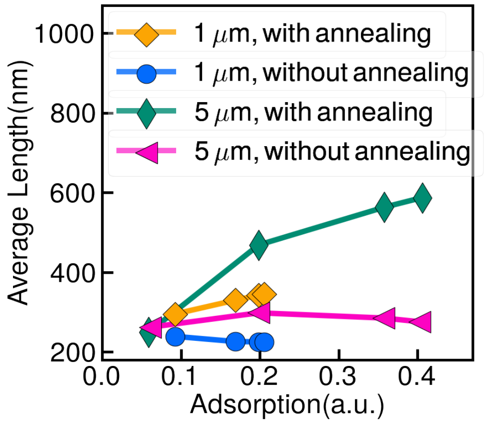


**Fig. S.15. Predictions of the average length vs adsorption at steady sate of mono-dispersed case.** The different colors and symbols denote different curvatures and with/without end-on annealing.

**H. Sensitivity analysis.** In addition to the inactivation analysis, we conducted a sensitivity analysis on these kinetic processes to examine how much the time-dependent adsorption varies according to each individual kinetic parameter. As the adsorption depends on multiple mechanisms, it is very time-consuming to conduct a sensitivity analysis across the entire parameters space. Therefore, we vary a single parameter while holding the others constant at their optimized value. The errors, defined as the relative difference of characteristic parameters, i.e., $T_{\mathrm{lag}}$, $S_{\mathrm{max}}$ and $n_{\mathrm{p}}$, against the corresponding values from experiments as follows:

$$ERR_T = \frac{1}{m}\sum_{i=1}^{m}|T_{\mathrm{lag}}/T^{\star}_{\mathrm{lag}} - 1|, \qquad ERR_S = \frac{1}{m}\sum_{i=1}^{m}|S_{\mathrm{max}}/S^{\star}_{\mathrm{max}} - 1|, \qquad ERR_n = \frac{1}{m}\sum_{i=1}^{m}|n_{\mathrm{p}}/n^{\star}_{\mathrm{p}} - 1|, \qquad \text{[S.9]}$$

in which $m = 4$ in mono-dispersed systems which represents the four different bulk concentrations for each bead assay and $m = 2$ in bi-dispersed case which represents the two curvatures.

The result of these sensitivity analysis are provided in Fig. S.20 to S.21. In all figures the left (A) and middle (B) compares the predictions of the characteristic parameters against the experimental results for 1 $\mu$m, for 5 $\mu$m mono-dispersed assays; the figure on the right (B) shows the same results and comparisons in the competitive (bi-dispersed) case. A brief description of the results are provided in the same page and subsection as the relevant figure.

***H.1. Varying cooperativity rate ($k_{on}^{coop}$).*** Comparing the errors in Fig. S.16 with the reference point where ($k_{\mathrm{coop}} = k^{\star}_{\mathrm{coop}}$), the errors are sensitive to changes in cooperativity, as seen in both mono-dispersed and bi-dispersed cases. Of particular note is the maximum slope, which the model predicts is dominated by the cooperativity mechanism, $J_{\mathrm{on}}^{\mathrm{coop}}$, and thus, shows more sensitivity when compared to $T_{\mathrm{lag}}$ and $n_{\mathrm{p}}$.

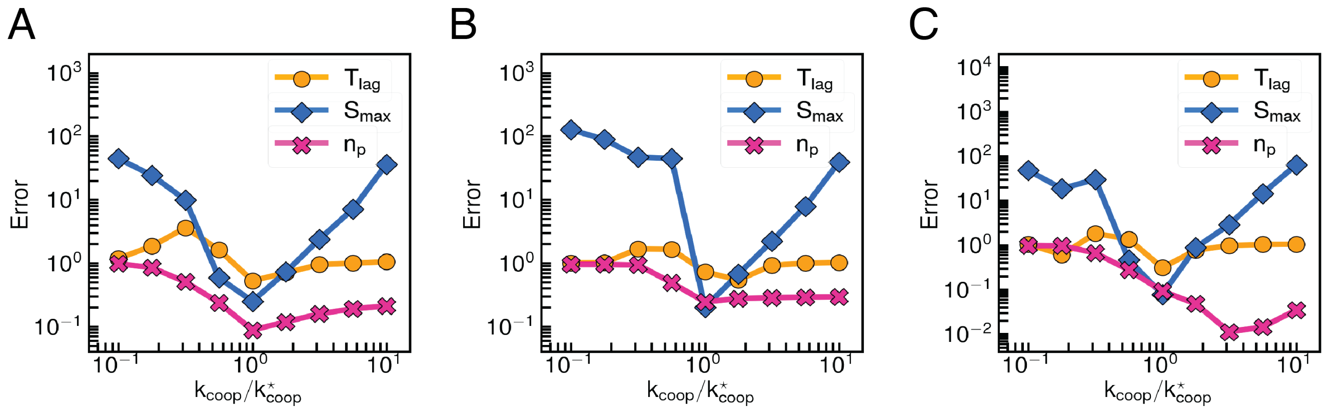


**Fig. S.16. Sensitivity analysis: cooperativity.** (A)mono-dispersed $1\ \mu$m beads, (B)mono-dispersed $5\ \mu$m beads, and (C)bi-dispersed assays.

***H.2. Varying fragmentation rate ($k_{frag}$).*** Comparing the errors in Fig. S.17 with the reference point where ($k_{\text{frag}} = k^{\star}_{\text{frag}}$) shows that errors are not a strong function of the fragmentation rate, especially the values of $T_{\text{lag}}$ and $n_{\text{p}}$. However, as was discussed in G.2, the length distribution can be altered significantly by the fragmentation rate. As the fragmentation rate approaches its extremes, filament length becomes either the maximum allowed value (too little fragmentation), or a single oligomer (too much fragmentation). Thus, to correctly capture length distribution the magnitude of the fragmentation rate should be comparable to the inverse of the time-scale of adsorption process in experiment, which is $\mathcal{O}(10^{-4})$ and consistent with our model prediction.

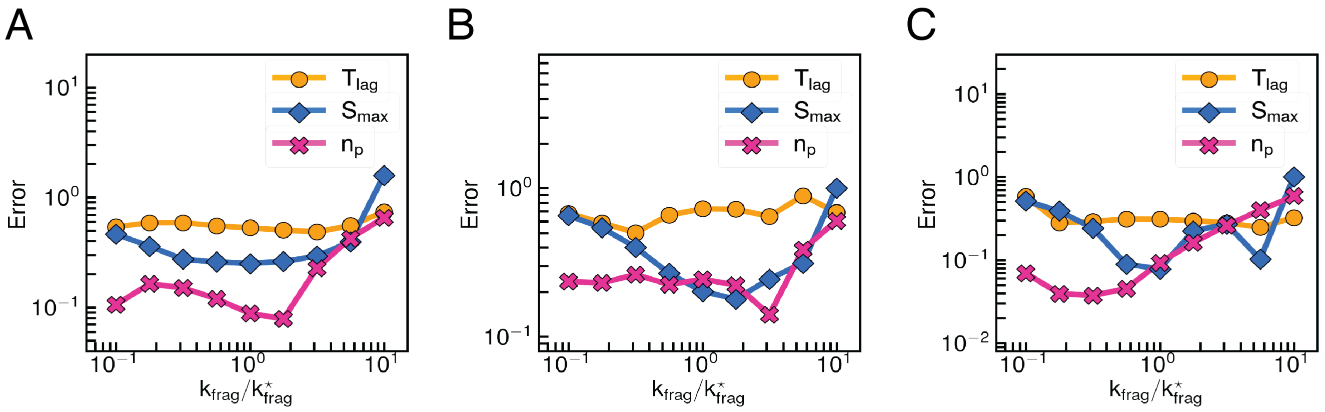


**Fig. S.17. Sensitivity analysis: fragmentation.** (A)mono-dispersed $1\ \mu$m beads, (B)mono-dispersed $5\ \mu$m beads, and (C)bi-dispersed assays.

***H.3. Varying defects dynamics ($\beta$).*** Comparing the errors in Fig. S.18 with the reference point where ($\beta = \beta^\star$) shows that both $T_{\rm lag}$ and $S_{\rm slope}$ are sensitive to changes in $\beta$, suggesting that the effective bulk concentration plays an important role in early adsorption. $n_{\rm p}$ is not a strong function of $\beta$, except at low $\beta$. This is because the adsorption plateau is independent of the bulk concentration when $n_{\rm b} \to \infty$.

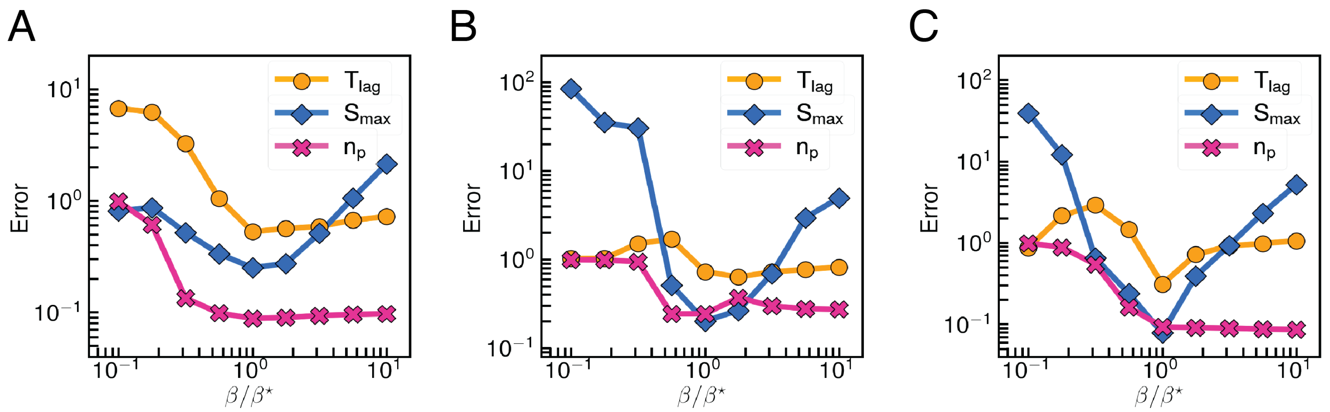


**Fig. S.18. Sensitivity analysis:** $\beta$**.** (A)mono-dispersed $1$ $\mu$m beads, (B)mono-dispersed $5$ $\mu$m beads, and (C)bi-dispersed assays.

***H.4. Varying surface density to adsorption ratio ($\Omega$).*** Comparing the errors in Fig. S.19 with the reference point where ($\Omega = \Omega^{\star}$) shows that all three characteristic parameters are sensitive to $\Omega$, especially $S_{\mathrm{slope}}$ and $n_{\mathrm{p}}$. This is a consequence of $\Omega$ controlling the conversion between light intensity and surface density and thus the predicted depletion magnitude. Either smaller or larger values of $\Omega$ result in underestimating or overestimating this depletion effect.

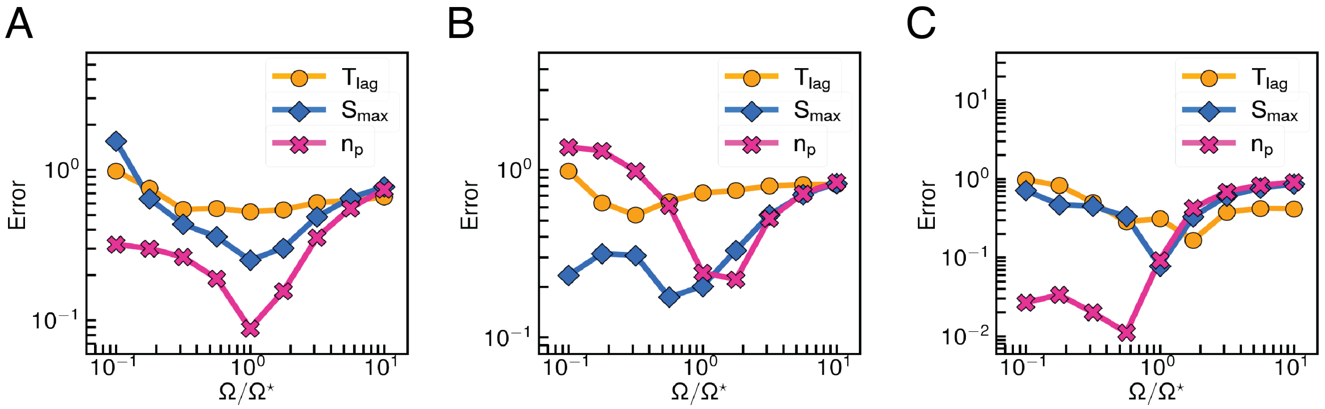


**Fig. S.19. Sensitivity analysis:** $\Omega$**.** (A)mono-dispersed $1\ \mu$m beads, (B)mono-dispersed $5\ \mu$m beads, and (C)bi-dispersed assays.

***H.5. Varying end-on annealing rate ($k_{anneal}$).*** Comparing the errors in Fig. S.20 with the reference point where ($k_{\text{anneal}} = k^{\star}_{\text{anneal}}$) shows that the errors are not a strong function of end-on annealing rate, which is consistent with our argument that annealing is not necessary to predict the adsorption. The errors of all the three parameters are in the same order of magnitude ($\mathcal{O}(10^{-1})$) as the reference point even the annealing rate changes a order of magnitude.

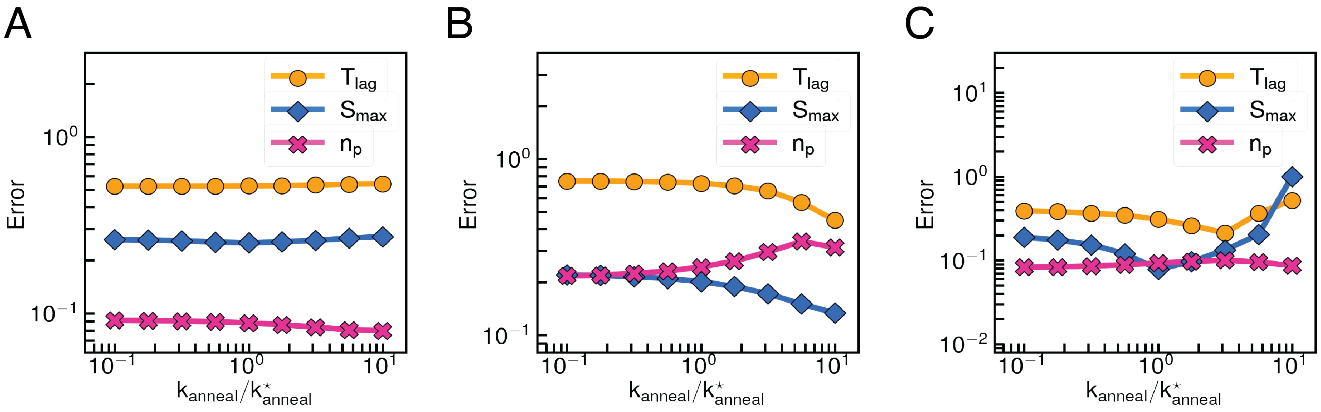


**Fig. S.20. Sensitivity analysis: annealing.** (A)mono-dispersed $1\ \mu$m beads, (B)mono-dispersed $5\ \mu$m beads, and (C)bi-dispersed assays.

***H.6. Varying effective rebinding exponent ($\xi$).*** Comparing the errors in Fig. S.21 with the reference point where ($\xi = \xi^\star$) shows that all three characteristic parameters are sensitive to $\xi$, especially $S_{\text{slope}}$ and $n_\text{p}$, as $\xi$ controls the stability (dwell time) of the filaments. Smaller values of $\xi$ result in a shorter dwell times, making it harder for adsorption to increase, while larger values of $\xi$ lead to the opposite behavior where adsorption increases too fast.

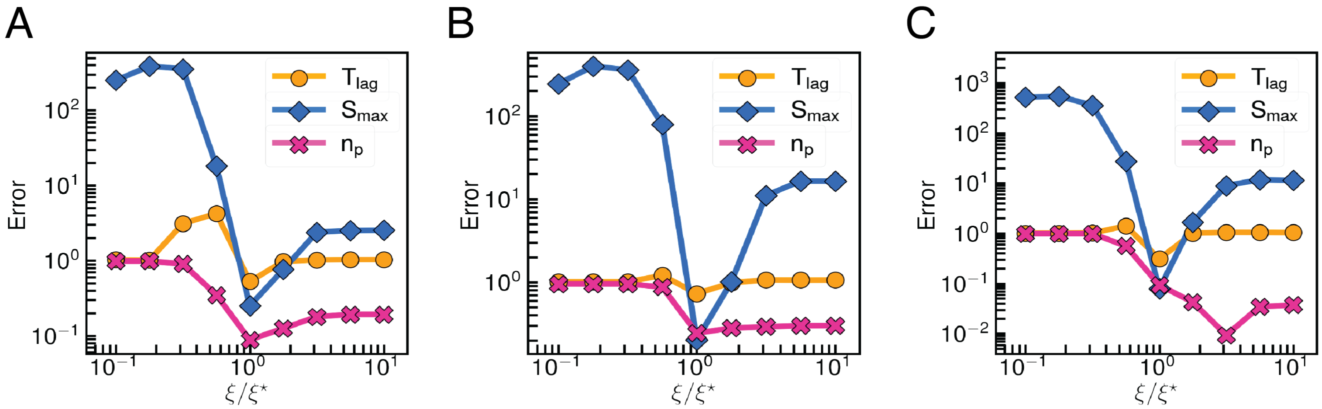


**Fig. S.21. Sensitivity analysis:** $\xi$**.** (A)mono-dispersed 1 $\mu$m beads, (B)mono-dispersed 5 $\mu$m beads, and (C)bi-dispersed assays.
